# Parent-of-origin effects in the UK Biobank

**DOI:** 10.1101/2021.11.03.467079

**Authors:** R. J. Hofmeister, S. Rubinacci, D. M. Ribeiro, Z. Kutalik, A. Buil, O. Delaneau

## Abstract

Identical genetic variations can have different phenotypic effects depending on their parent of origin (PofO). Yet, studies focussing on PofO effects have been largely limited in terms of sample size due to the need of parental genomes or known genealogies. Here, we used a novel probabilistic approach to infer PofO of individual alleles in the UK Biobank that does not require parental genomes nor prior knowledge of genealogy. Our model uses Identity-By-Descent (IBD) sharing with second- and third-degree relatives to assign alleles to parental groups and leverages chromosome X data in males to distinguish maternal from paternal groups. When combined with robust haplotype inference and haploid imputation, this allowed us to infer the PofO at 5.4 million variants genome-wide for 26,393 UK Biobank individuals. We used this large dataset to systematically screen 59 biomarkers and 38 anthropomorphic phenotypes for PofO effects and discovered 101 significant associations, demonstrating that this type of effects contributes to the genetics of complex traits. Notably, we retrieved well known PofO effects, such as the *MEG3/DLK1* locus on platelet count, and we discovered many new ones at loci often unsuspected of being imprinted and, in some cases, previously thought to harbour additive associations.

## Introduction

Parent-of-Origin (PofO) effects refer to genetic variations having an effect on a phenotype that depends on the parent from which alleles are inherited^1,2^. PofO effects are thought to mainly result from genomic imprinting, a mechanism relying on parent-specific DNA methylation, named imprints, that silence one of the parental copies of a gene. Such parent-specific imprints are established during spermatogenesis and oogenesis and are maintained in all somatic cells of the offspring^3^. This leads to some genes, called imprinted genes, to exhibit an allele-specific expression pattern that depends on the PofO of the underlying genetic sequence. This allele-specific expression can be maintained throughout life or specific to some development states^4^. One of most studied imprinted loci in the human genome is the H19 loci at 11p15.5 that is involved in growth and development disorders such as the Beckwith–Wiedemann or Silver–Russel syndromes^5^. Multiple studies have investigated PofO effects on complex traits, notably for the *KCNQ1* and *KLF14* genes whose associations with type 2 diabetes risk depends only on the maternal copies^6^, and the well-known *MEG3/DLK1* imprinted locus associated with age at menarche^7^ and platelet count^8^.

Searching for PofO effects on a genome-wide scale requires knowing the PofO of each individual allele. The most direct approach to obtain this information relies on the availability of parental genomes, which allows using the Mendelian principles of inheritance to determine the parent from which a specific allele is inherited. Study cohorts usually include a small number of genotyped parent-offspring duos and trios, resulting in a low discovery power and a challenging detection of PofO effects. To alleviate this problem, multiple approaches have been explored so far. First, by deploying large efforts in data collection, such as the study performed on the DiscovEHR cohort^9^, representing the largest PofO study done to date, by regrouping more than 22,000 samples with at least one genotyped parent and hundreds of phenotypes assessed. Alternatively, this can also be achieved by meta-analysis across multiple cohorts regrouping duos and trios, with the caveat of restricting the analysis to the subset of phenotypes in common across datasets^7,10^. Second, statistical approaches have been proposed to test for PofO effects in large collections of unrelated samples by exploiting the differences in phenotypic variance between heterozygous and homozygous individuals, with the caveat of also detecting effects unrelated to PofO such as gene-environment interactions^11^. Third, it has been shown that the PofO of an individual’s alleles can also be determined by the use of cousins as *surrogate parents* when parental genomes are not available^6^. This latter approach is particularly well suited for datasets comprising many samples from the same generation but also requires the genealogy of all individuals in the study cohort, which is not the case in large datasets such as the UK Biobank^12^.

In this work, we present a probabilistic method to infer the PofO of alleles in biobank scale datasets from second- and third-degree relatives without requiring any parental genomes nor explicit genealogy to be known. To do so, our approach combines multiple estimation steps, involving surrogate parent group formation, parental status assignment based on chromosome X, haplotype inference, Identity-By-Descent (IBD) detection and haploid imputation. When applied to the UK Biobank dataset, this allows us to infer the PofO for 21,484 samples with high confidence in addition to the 4,909 duos/trios for which we could perform direct inference from parental genomes, resulting in a dataset comprising a total of 26,393 samples. Using duos/trios as ground truth, we show that our PofO estimations from second- and third-degree relatives have a high call rate (~75%) and low error rate (<1%) at heterozygous genotypes. Taking advantage of the vast phenotypic diversity of the UK Biobank, we carried out genome-wide association scans for PofO effects for 97 phenotypes, retrieving the effects of well-known imprinted loci as well as discovering novel PofO associations, demonstrating the high discovery power of our method and revealing the diversity and complexity of PofO effects in the human genome. All the summary statistics for the conducted association scans are publicly available online (http://poedb.dcsr.unil.ch/).

## Results

### PofO inference from genotype data

To infer the PofO of all alleles carried by a given target sample, we proceed in two consecutive steps as detailed below:

1. *Identification of surrogate parents* (**Figure 1A**). For each target sample, we identify close relatives and we determine which of the two parents (mother or father) conveys the relatedness. For this, we first look at pairwise kinship estimates given by KING^13^ to identify second or third degree relatives and group them into the two parental groups based on their relatedness: they cluster in the same group if the samples are related and in different groups otherwise. Then, we assign parental status (maternal or paternal) to parental groups for male targets only by exploiting the fact that their single chromosome X copy is maternally inherited. Therefore, we search for relatives sharing portions of their chromosome X Identical-By-Descent (IBD) with the target and we label them as surrogate mothers. We also propagate the information to other relatives: those from the same parental group are also labelled as surrogate mothers and those from the other parental group as surrogate fathers. In case no IBD is found, we cannot annotate parental groups as maternal or paternal and we exclude the target sample from the dataset. Hereafter, we call *surrogate parents* the close relatives we identified using this approach.
2. *Assignment of PofO to alleles* (**Figure 1B**). After the identification of surrogate parents, we assign PofO to the target’s alleles. First, we search for autosomal DNA segments shared IBD between the target and the surrogate parents using IBD mapping robust to both phasing and genotyping errors (see Methods, **Figure S1**). Then, we classify the resulting IBD segments as being maternally or paternally inherited depending on the surrogate parent they map to. This delimits a subset of alleles that are co-inherited from the same parent within and across chromosomes (i.e. that co-localize on the transmitted set of homologous chromosomes). This leaves another subset of alleles for which we do not know the PofO (i.e. those not shared IBD with any of the surrogate parents). For those, we extrapolate the PofO using statistical phasing: we model alleles for which we know the PofO status as a haplotype scaffold^14^ onto which all remaining alleles are probabilistically phased using SHAPEIT4^15^ (**Figure S1**). The PofO assignment of these remaining alleles is then given by their frequency of co-localization onto each haplotype scaffold, which also reflects how reliable the phasing is (i.e. phasing certainty, **Figure S1**). Finally, we extrapolate the PofO for untyped variants by performing haploid imputation of each parental haplotype in turn using IMPUTE5^16^ and the HRC as reference panel^17^.

**Figure 1:**
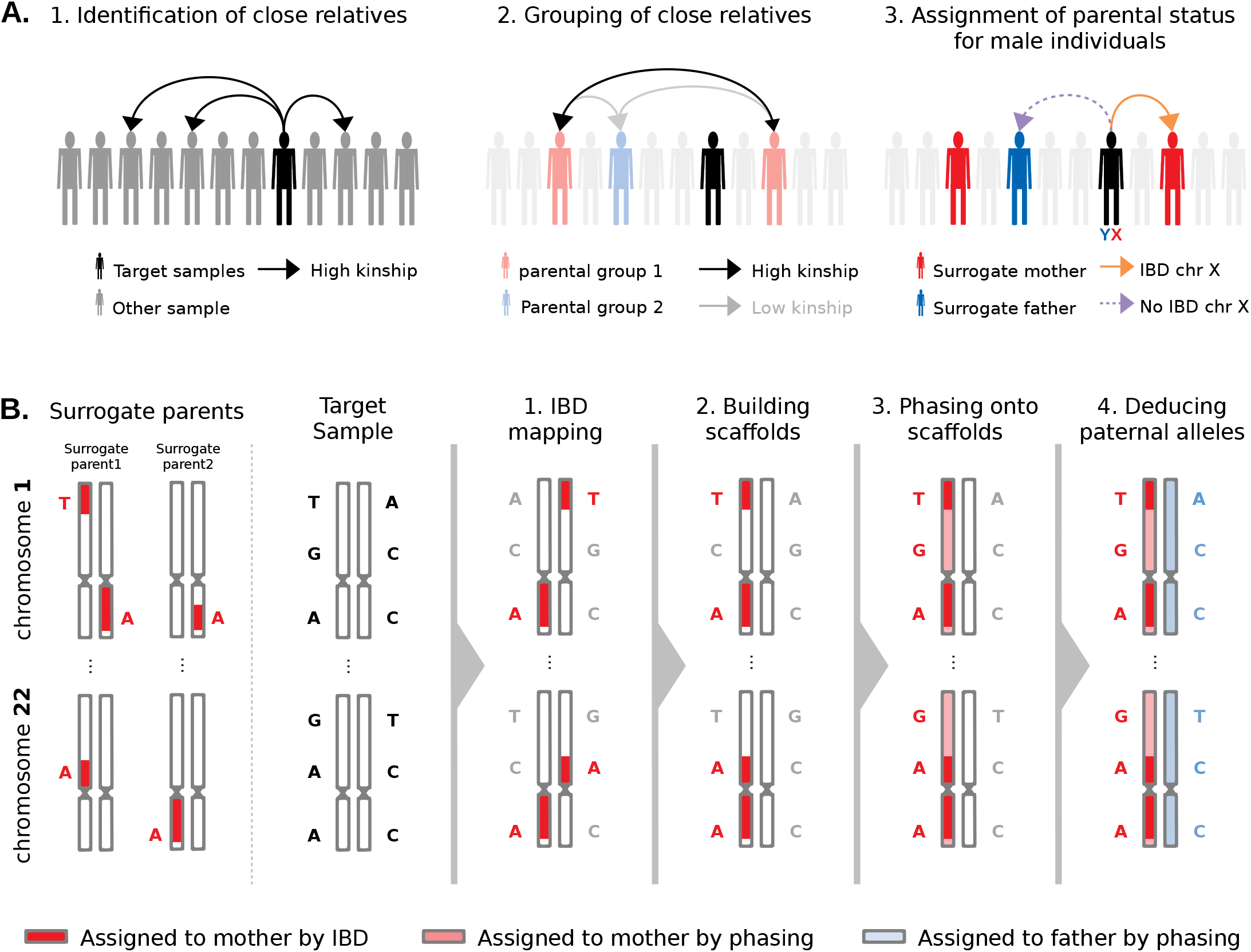
Rationale of PofO inference. **A.** Identification of surrogate parents in 3 steps: (1) identification of close relatives for a target sample of interest using the pairwise kinship estimates, (2) clustering of close relatives by maximizing and minimizing the inter- and intra-groups, respectively, (3) assignment of parental status to close relatives’ groups (i.e. surrogate parents) using IBD sharing on chromosome X for male targets. **B.** Parent-of-origin inference in 4 steps: (1) identification of autosomal IBD segments shared between the target and the surrogate parents, (2) scaffold construction with co-inherited alleles localized on the same homologous chromosome across all autosomes, (3) statistical phasing of all remaining alleles against the scaffold and (4) whole genome deduction of the maternal and paternal origins of alleles from phasing probabilities.

### Validation of the PofO inference on duos and trios

To assess the accuracy of our approach, we used 443,993 genotyped UK Biobank samples from British and Irish ancestry together with their pairwise kinship estimates in order to identify a subset of samples with parents and second-to-third degree relatives. For these samples, we inferred the PofO using two approaches: directly from the parents or using second-to-third degree relatives as surrogate parents. We compared the quality of the PofO inference given by surrogate parents to the direct approach based on parental genomes that we consider to be the ground truth. We found a total of 3,872 parent-offspring duos and 1,037 trios, of which 1,090 duos and 309 trios also have groups of surrogate parents. We used this subset of 1,399 samples to assess optimal parameters and accuracy of the method. We focussed on two metrics: (i) the error rate, which is the percentage of heterozygous genotypes with incorrect PofO assignment and (ii) the call rate, which is the percentage of heterozygous genotypes for which a PofO call could be made (see Methods). We explored a range of different parameter settings for the IBD detection and PofO confidence score (i.e. phasing certainty onto haplotype scaffold). We empirically found that using haplotype segments longer than 3cM as scaffold and a phasing certainty above 0.7 lead to a good trade-off between call rate and error rate (**Figure 2A**). This resulted in a whole genome error rate and call rate of 0.51% and 74.5%, respectively. As expected, the error and call rate depend on the number of available surrogate parents per target (**Figure 2B**), with the call rate increasing and the error rate decreasing as the number of surrogate parents increases. The majority of our targets have only a single surrogate parent (75.95% of the target samples, **Figure 2C**) and even in this case, a call rate of 70.9% and an error rate of 0.6% is achieved (**Figure 2B**). We then considered the genomic localization of variants: we found a lower call rate and a slightly higher error rate as we approach telomeres, which results from phasing edge effects (**Figure 2D**). Overall, we found that the vast majority of variants have a low error rate: 56% of variants are inferred perfectly and 79% have an error rate <1% (**Figure 2E**). This low error rate mostly results from the high phasing accuracy that can be achieved in the UK Biobank using SHAPEIT4^15^. Overall, we obtained a whole genome switch error rate of 0.0845% between consecutive heterozygous genotypes when comparing to parental genomes, with only small variations across chromosomes (**Figure S2A**). When looking at the distribution of these switch errors along the genome, we found that they mostly occur within small segments and that long range errors are almost entirely corrected by the use of haplotype scaffolds (**Figure S2B**). As a result, we obtained haplotypes that are resolved across entire chromosomes with only a few sporadic errors that we believe to mostly result from genotyping errors given their frequency (<0.1% error rate).

**Figure 2:**
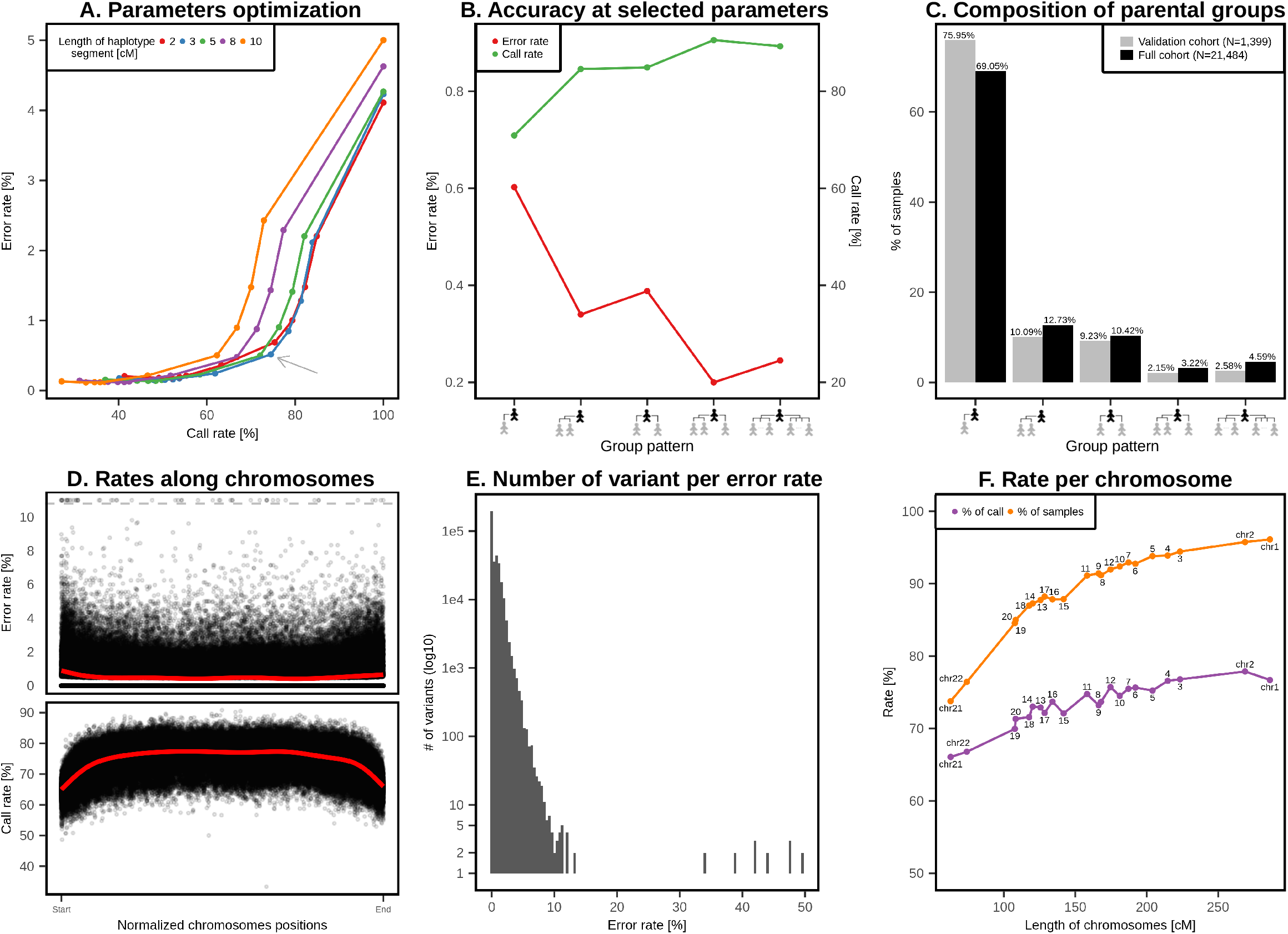
Validation of the PofO inference. **A.** Call rate (x-axis) and error rate (y-axis) as a function of (i) the minimal length of IBD tracks for scaffold construction and (ii) the minimal phasing probability used to call a heterozygote as phased. Each point corresponds to a given phasing probability threshold going from 0.5 (right most point) to 1.0 (left most point) with steps of 0.05. The black arrow indicates the parameters we used in our analysis (3cM long IBD tracks and 0.7 minimal phasing probability). **B.** Call rate (left y-axis) and error rate (right y-axis) as a function of the composition of the parental groups (x-axis). The latter ranges from one parental group with one surrogate parent (left) to two parental groups comprising multiple surrogate parents (right). **C.** Fraction of targets as a function of the composition of the parental groups (x-axis): in the validation data (N=1,399) in grey and in the call set (N=21,484) in black. **D.** Error rate per variant site (top panel; y-axis) and call rate (bottom panel; y-axis) as a function of their normalized positions relative to each telomere (x-axes). Red lines are fitted density curves. Error rates greater than 10% are capped to 11% as indicated by the dashed grey line. **E.** Distribution of error rates per number of variant sites (y-axis is on the log scale). **F.** Fraction of samples (purple) and heterozygotes (i.e. call rate; orange) in the call set for which PofO is inferred, as a function of chromosome length (cM, x-axis). Chromosome numbers are shown next to the points in grey.

### PofO inference in more than 25,000 individuals

For all genotyped British and Irish samples in the UK Biobank without any genotyped parent (N=438,993), we inferred the PofO using the method based on surrogate parents, as described above. In total, we found 105,826 samples with second to third degree relatives forming groups of surrogate parents. Amongst those, we assigned parental status to surrogate parent groups for a subset of 21,484 samples using IBD matching on chromosome X. Comparing the distribution of surrogate parents per target sample, we found a remarkable match between the full (N=21,484) and the validation (N=1,399) datasets (**Figure 2C**), suggesting that we can expect similar error rates between datasets.

As our method requires IBD sharing between the targets and the surrogate parents, no inference can be made for chromosomes where no IBD sharing is found. The number of samples with PofO inference thus varies across chromosomes, ranging from 15,645 samples (72.8%) for chromosome 21 to 20,381 samples for chromosome 1 (94.8%), depending on chromosomal length (**Figure 2F**). It follows that the call rate also varies across chromosomes, ranging from 66% for chromosome 21 to 77.9% for chromosome 2 (**Figure 2F**). From the sample point-of-view, we found that 31.3% of the samples have inference for the 22 autosomes and 96.1% have inference for more than 15 chromosomes (**Figure S3**).

Finally, we merged the 21,484 samples with PofO inferred from surrogate parents together with the 4,909 samples with PofO inferred from genotyped parents (3,872 duos and 1,037 trios) to get a final set comprising a total of 26,393 UK Biobank individuals with PofO inference across 5.4 million variants genome-wide. Together with deep phenotyping, this dataset represents a unique data set to study PofO effects on complex traits.

### PofO effects contribute to complex traits

In this study, we used four association models: (i) *maternal*, to test maternally inherited alleles, (ii) *paternal,* to test paternally inherited alleles, (iii) *differential,* to compare paternally and maternally inherited alleles at heterozygous genotypes only and (iv) *additive,* as a control to test minor alleles regardless of PofO. We ran a GWAS scan for each model using BOLT-LMM^18^ on 97 quantitative phenotypes of the UK biobank (**Table S1**). **Figure 3A** shows an example of these 4 types of GWAS scans for platelet count. Summary statistics for all 396 GWAS scans are publicly available online (http://poedb.dcsr.unil.ch/).

**Figure 3:**
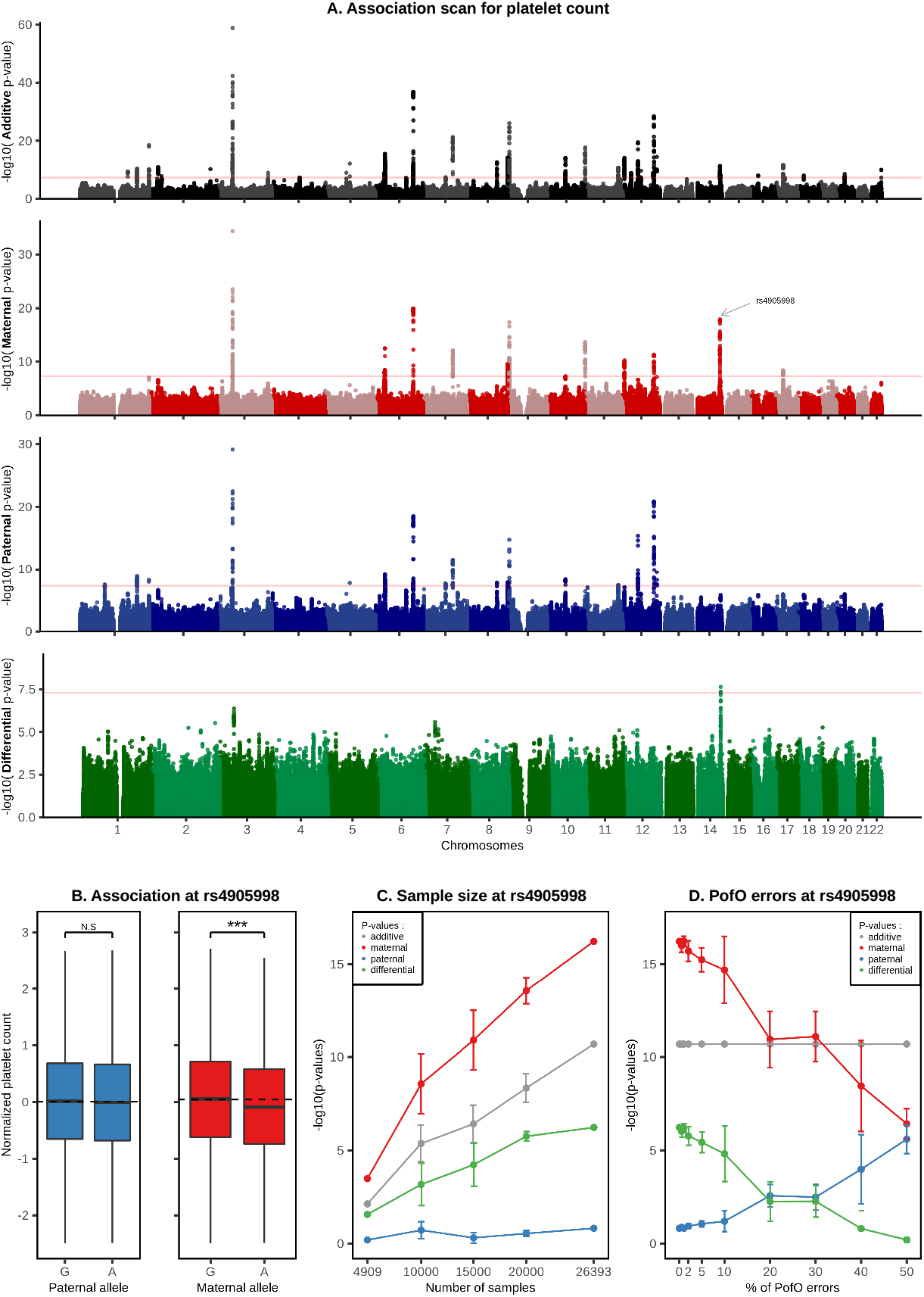
Association scans for PofO effects on platelet count. **A.** Manhattan plots of four association scans with platelet count. From top to bottom plots are shown results for additive (black), maternal (red), paternal (blue) and differential (green) models. **B.** Box plot of the normalized platelet counts (y-axis) stratified by risk alleles and origin at SNP rs4095998; paternal in blue and maternal in red (x-axis). The horizontal dotted lines represent the phenotypic median of the risk allele A. **C.** Association strength as -log10(p-value) for rs4905998 (y-axis) as a function of the number of randomly chosen samples included in the analysis under the additive (black), paternal (blue), maternal (red) and differential (green) models. Each point for N=[10,000; 15,000; 20,000] represents the median p-value obtained after 10 randomizations with vertical bars representing the standard error. Points for N=4,909 and N=26,393 represent the p-values obtained using only the samples with genotyped parents and using our full sample size, respectively. **D.** Association strength as – log10(p-value) for rs4905998 (y-axis) as a function of the fraction of samples in the full call set for which PofO has been randomly drawn (x-axis). Each point represents the median p-value obtained after 10 randomizations with vertical bars representing the standard errors.

Given that multiple association scans are performed per phenotype, we used a two-step approach to find significant PofO associations. In a first pass, we selected all associations directly genome-wide significant (Bonferroni; p-value < 5×10^-8^) in the differential association scan and found 9 of them. This pass is primarily designed to capture PofO effects exhibiting strong opposite effects between the two parents; effects that do not necessarily reach genome-wide significance in the paternal or maternal scans. In a second pass, we defined a set of 631 candidate associations by selecting all associations genome-wide significant either in the maternal scan, in the paternal scan, or simultaneously in both scans (**Figure S4A)**. Then, we distinguished PofO effects from additive effects by extracting associations significant at 1% FDR in the differential scan. We found 92 of these associations **(Figure S4B-D)**. Conversely to the first pass, this second pass is primarily designed to capture PofO effects with only one strong parental effect (either maternal or paternal); effects that do not necessarily reach genome-wide significance in the differential scan. Of note, all association loci we report have been trimmed to only retain one candidate SNP per locus (see method).

Overall, we scanned 97 quantitative phenotypes from 2 phenotypic categories: 59 biomarkers (28 blood chemistry, 31 blood count) and 38 anthropomorphic traits (32 body composition by impedance and 6 body size measurements) (**Table S1**). We discovered 101 significant PofO associations for 60.8% (n=59) of the tested phenotypes: 45 for biomarkers and 56 for anthropomorphic traits (**Table S2**). These associations are widespread throughout the genome (**Figure 4**), suggesting that PofO effects are more common in complex traits than previously thought. We found no significant enrichment towards a specific parental origin either across all phenotypes taken together (47 maternal effects versus 46 paternal effects) or across phenotypic categories (**Table S3**).

**Figure 4:**
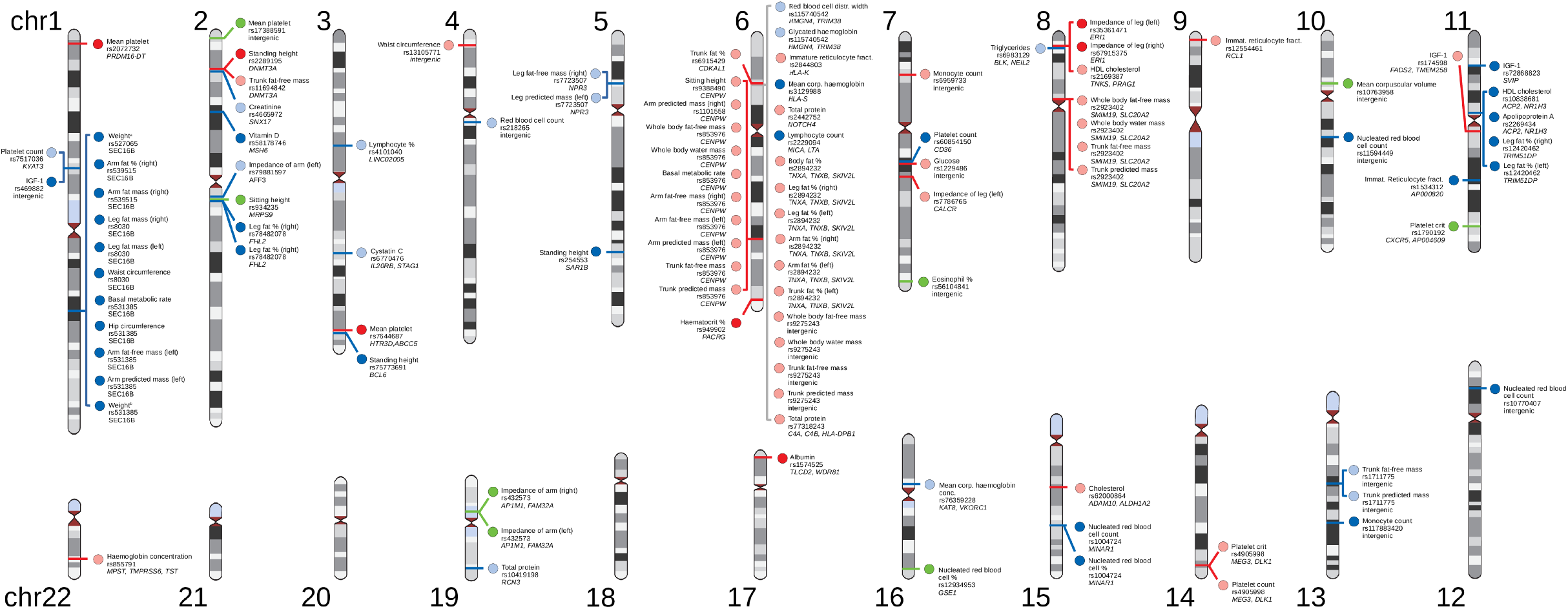
PofO effects in the human genome. Genomic locations of the 101 significant PofO associations discovered in this study. Each dot indicates a PofO signal with the phenotype label, the variant site id and the mapped gene, if any, are shown next to it. Maternal, paternal and differential signals are shown in red, blue and green, respectively. Paternal and maternal effects with positive effect size are indicated by dark colours (i.e. dark blue and dark red, respectively). Paternal and maternal effects with negative effect size are indicated by light colours (i.e. light blue and light red, respectively).

Then, we tested the 101 PofO significant associations for additive effects in the exact same sample set and found that 84 do not reach genome-wide significance (**Table S2**). For these, we found the additive p-values strongly anti-correlated with the differential p-values (Pearson R= −0.92, p-value = 7.7×10^-35^, **Figure S5**). This likely occurs because the opposite effects of maternal and paternal alleles are cancelled out in the additive model which considers these alleles together. When scaling to the full UK Biobank sample collection (N=420,531 Europeans, Pan-UKB team. https://pan.ukbb.broadinstitute.org), 31 associations still remain non-significant, suggesting that these associations can only be detected when considering PofO information. Conversely, it also means that the underlying genetic effect of 70 associations reported as additive effects in the UK Biobank is of parental origin, suggesting that many additive associations reported so far in GWAS capture PofO effects.

### Replication of known PofO associations

PofO effects are mostly attributed to genomic imprinting^1,2^. For instance, in the largest PofO study to date, 10 significant associations have been reported within imprinted loci^9^. Out of these 10 associations, we could assess 7 of them in our dataset (the same SNP-phenotype pair exists in UK Biobank) and could replicate 5 as PofO signals with the exact same parent and direction of effects (**Table 1**).

**Table 1:** Replication of PofO associations from Kim et al.^9^.

| Phenotype-SNP pairs |  |  | DiscovEHR cohort (Kim et al.) |  |  |  |  |  |  |  |  | UK Biobank cohort |  |  |  |  |  |  |  |
| --- | --- | --- | --- | --- | --- | --- | --- | --- | --- | --- | --- | --- | --- | --- | --- | --- | --- | --- | --- |
| Phenotype | SNP | Risk allele | N (Add) | N (PofO) | MAF | Add. P-value | Add. Beta | Pat. P-value | Pat. Beta | Mat. P-value | Mat. Beta | N (PofO & Add.) | MAF | Add. P-value | Add. Beta | Pat. P-value | Pat. Beta | Mat. P-value | Mat. Beta |
| % Monocytes | rs117989553 | C | 102,797 | 17544 | 0.055 | 9.6E-27 | -0.09 | 1.9E-05 | -0.13 | 0.15 | -0.04 | 24454 | 0.0598 | 3.9E-10 | -0.113 | 1.6E-04 | -0.096 | 5.6E-06 | -0.126 |
| % Monocytes | rs117421106 | T | 102,797 | 17544 | 0.023 | 3.2E-19 | -0.12 | 7.9E-05 | -0.19 | 0.084 | -0.08 | 24454 | 0.0266 | 4.7E-04 | -0.089 | 1.7E-04 | -0.138 | 0.27 | -0.05 |
| Cholesterol | rs117515500 | T | 93,077 | 14619 | 0.031 | 1.1E-07 | 0.07 | 0.14 | 0.07 | 1.5E-05 | 0.2 | 24074 | 0.0344 | 0.66 | -0.011 | 0.74 | -0.009 | 0.67 | -0.026 |
| HDL-C | rs12154627 | C | 93,38 | 14642 | 0.481 | 1.2E-23 | 0.04 | 0.064 | 0.03 | 2.1E-16 | 0.13 | 22109 | 0.4917 | 0.0018 | 0.031 | 0.6 | 0.001 | 2.7E-07 | 0.066 |
| Triglyceride | rs12154627 | C | 93,365 | 14625 | 0.481 | 5.1E-17 | -0.04 | 0.56 | -0.01 | 1.4E-09 | -0.1 | 24051 | 0.4917 | 5.7E-04 | -0.035 | 0.42 | -0.015 | 1.6E-05 | -0.06 |
| Bilirubin | rs4758459 | T | 108,535 | 17913 | 0.2 | 8.2E-09 | 0.03 | 0.073 | 0.03 | 5.3E-05 | 0.07 | 23975 | 0.2022 | 0.061 | 0.014 | 0.89 | 0.001 | 3.5E-04 | 0.044 |
| Platelet crit | rs10146962 | C | 115,308 | 19552 | 0.329 | 3.2E-20 | -0.04 | 0.12 | 0.02 | 2.3E-11 | -0.1 | 24493 | 0.3387 | 1.5E-11 | -0.062 | 0.14 | -0.021 | 7E-17 | -0.112 |

The most notable replicated hit relates to the *MEG3/DLK1* locus at which SNP rs10146962 is associated with platelet crit (**Table 1**). This association has been previously described for another SNP in the same locus: SNP rs1555405 associated with decreased platelet count only when the A allele is maternally inherited^8^. In our study, we replicated these two associations: rs10146962 (maternal p-value=7×10^-17^, paternal p-value=0.14, **Table 1**) and rs1555405 (maternal p-value=2×10^-16^, paternal p-value=0.13). However, we found a stronger association at the same locus for rs4905998 (**Figure 3A-B**, **Table S2 hit #90**), a SNP in high linkage disequilibrium with both rs10146962 (R^2^=0.87) and rs1555405 (R^2^=0.72) and eQTL for *MEG3* in whole blood (p-value = 6.5×10^-5^; https://gtexportal.org). This variant is associated with platelet count in the additive, maternal and differential scans (p-values = 6.2×10^-12^, 1.2×10^-18^ and 4.9×10^-8^), but not in the paternal scan (p-value=0.18), which confirms the strong maternal effect at this locus while providing a new promising candidate causal variant (**Figure 3A-B**).

We then investigated the benefit of using our approach on the discovery power using this specific association as an illustration. To do so, we used 4,909 UK Biobank duos/trios as baseline and gradually added random subsets of 5,000 samples for which PofO inference could be made from surrogate parents, ending up with the full set of 26,393 samples. Increasing the sample size led to a clear boost in association strength for the additive, maternal and differential signals, with a maternal scan ranging from weak significance on the duos/trios (n = 4,909; p-value = 3.27×10^-04^) to strong significance on the full sample size (n = 26,393; p-value = 1.2×10^-18^; **Figure 3C**), while the paternal signal remained largely non-significant. Similarly, we also looked at the effects of errors in the PofO inference on the discovery power by randomizing the PofO assignment for an increasing number of samples. As expected, this progressively diluted the maternal signal onto the two parental origins while leaving the additive signal unchanged (**Figure 3D**).

Another well-established PofO association relates to four SNPs (rs2237892, rs231362, rs2334499 and rs4731702), falling in two distinct regions that harbour well-documented imprinted gene clusters, 11p15.5^19^ and 7q32^20,21^, that have been found to exhibit PofO effects on type 2 diabetes^6^ (T2D). As we could not directly test T2D status because of insufficient sample size, we instead investigated associations of these four SNPs with all the 97 tested phenotypes and found multiple associations (differential p-values < 0.01; **Table 2**). For example, we found rs4731702 at 7q32 with opposite parental effects on the lymphocyte-neutrophil ratio, previously proposed as a potential prognostic biomarker for T2D^22–24^. The same SNP also shows maternal effects on HDL cholesterol and apolipoprotein A, both tightly related to T2D^25–28^. At 11p15.5, we found rs2237892, rs231362 and rs2334499 associated through different types of PofO effects with Cystatin C, another putative biomarker for T2D^29–31^. Finally, the T allele of rs2334499 seems to decrease multiple weight- and fat-related phenotypes when paternally inherited, and conversely increase them when maternally inherited. Overall, we found 22 associations between these four SNPs and phenotypes that closely relate to T2D status, thereby providing hypotheses of the impact of these T2D risk alleles on other relevant phenotypic layers.

**Table 2:**
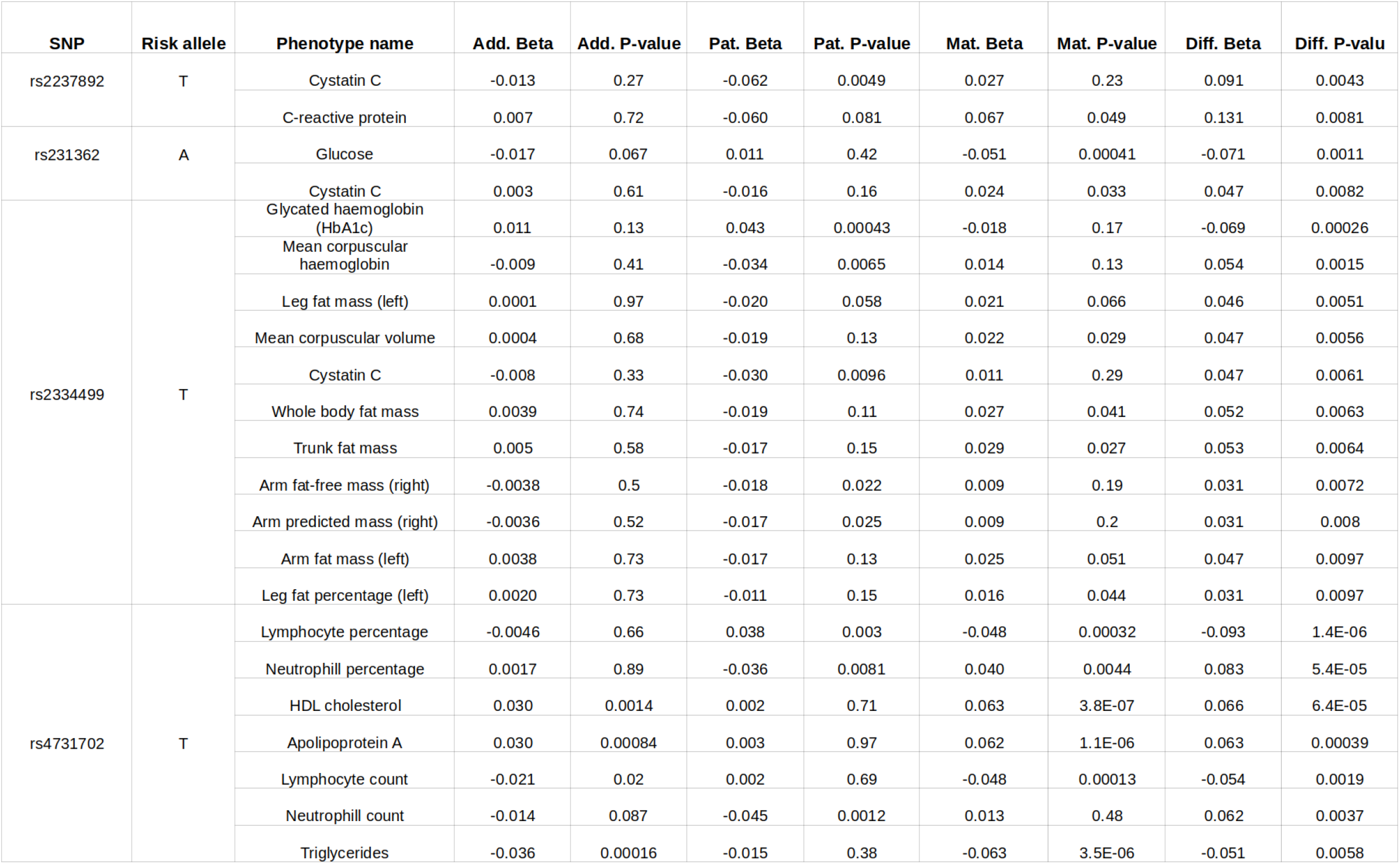
Replication of PofO associations from Kong et al.^6^.

### Discovery of novel PofO associations

The vast majority of the PofO associations we discovered in this study have not yet been reported in the literature and only 3 of them could be directly linked to well-documented imprinted genes^32^ (see URLs; **hits #65,90,91**). This study therefore represents a valuable resource to further characterize PofO effects and to provide additional candidate imprinting genes in the human genome. As a proof-of-principle, we highlight in the following four key examples of significant associations bringing biological insights of interest on PofO effects and genomic imprinting.

A first example is the maternal association between rs7786765 at 7q21.3 and leg impedance, a measurement used to estimate body fat and muscle mass (maternal p-value=2.1×10^-8^, paternal p-value=0.065; **Table S2 hit #65**). This SNP is intronic to *CALCR* (calcitonin receptor), a gene catalogued as maternally expressed^32^ (see URLs). Supporting this discovery, the gene encoding calcitonin itself *(CALCA)* has been associated with phenotypes related to leg impedance, such as body weight and body fat^33^. This represents an expected result: a PofO effect on a gene, relevant to the tested phenotype, and known to be imprinted with a parental expression consistent with the association.

A second example is the maternal association between rs12554461 at 9p24.1 and immature reticulocyte fraction (maternal p-value = 2×10^-8^, paternal p-value = 0.066; **Table S2 hit #75**). This SNP is intronic to *RCL1* and affects its expression across multiple brain tissues (p-values < 5.0×10^-5^; https://gtexportal.org). Interestingly, *RCL1* has been described to be differentially methylated in an imprinted manner^34^ but never fully characterized as imprinted. In addition, no clear evidence of PofO effects have been reported so far for *RCL1,* although it has been repeatedly associated with blood cell phenotypes^35,36^. Here, the PofO signal for *RCL1* brings additional evidence of its involvement in blood cell phenotypes and of its imprinted status with maternal expression.

A third example is the paternal associations between four SNPs in high LD (R^2^ > 0.5 between rs527065, rs539515, rs8030 and rs531385) with weight, waist circumference, hip circumference, basal metabolic rate and arm/leg mass (paternal p-values < 5×10^-8^, maternal p-values > 0.05; **Table S2 hits #4-14**). All these SNPs locate in the locus 1q25.2, most of them being either intronic or splicing QTLs for *SEC16B* and repeatedly found to be associated under an additive model with multiple weight- or obesity-related phenotypes^37–39^. Our data suggests that a paternal effect underlies this well-known association with the downstream gene, *SEC16B,* being a new imprinted candidate with paternal expression.

A last example is the maternal association between rs2289195 at 2p23.3 with standing height (maternal p-value = 5.5e^-10^, paternal p-value = 0.035; **Table S2 hit #16**). This SNP is intronic to *DNMT3A,* a gene playing a key role in the formation of imprints during gametogenesis^40–42^. In addition, mutations within *DNMT3A* have been associated with an overgrowth syndrome, characterised among others by an increased height^43^. Taken together, this suggests a possible mechanism in which variations in 2p23.3 alter *DNMT3A* function which results in the perturbation of maternal imprints formation and ultimately in increased standing height.

## Discussion

Studying PofO effects requires parental genomes or genealogies to determine the set of alleles transmitted by each of the two parents to the offspring. As a consequence, studies focussing on this type of effects usually rely on relatively small sample sizes and are therefore underpowered to discover genetic effects as those typically involved in complex traits. In this work, we propose a novel approach that leverages the high degree of relatedness between individuals inherent to biobank-scale datasets in order to infer the PofO of alleles for many individuals and variant sites without any parental genomes or genealogy being available. When applied on UK Biobank, this approach could predict the PofO of alleles for around 5% of the total number of samples, resulting in a dataset comprising the PofO inference for more than 26,000 samples at 5.4 million variants. Together with deep phenotyping, this dataset allows studying PofO effects on a larger scale than previous studies and considerably increases discovery power, as demonstrated by our ability to replicate known associations as well as discover new ones.

Overall, we tested 97 phenotypes and found evidence of PofO effects for 59 of them, suggesting that this type of genetic effect is commonly involved in complex traits. Out of the 101 distinct associations we mapped, the strongest one is a maternal effect on platelet count/crit located in the *MEG3/DLK1* imprinted locus; an association that has already been described in multiple studies^8,9^. Beyond mapping PofO effects at known imprinted genes (e.g. *MEG3, CALCR),* we also found PofO effects at genes for which imprinting has not yet been clearly established (e.g. *RCL1)* or not even suspected (e.g. *SEC16B),* suggesting that the current annotation of imprinted genes is still far from being complete or that the molecular mechanism underlying PofO effects is not necessarily directly linked to genomic imprinting^44^. Overall, we believe that 101 PofO effects we reported in this work is only the emerging part of the iceberg and many other associations could be revealed by meta-analysis summary statistics of this study (http://poedb.dcsr.unil.ch/) together with those from independent studies (e.g. *KLF14*).

One of the strengths of our approach resides in its ability to make PofO calls with a low error rate. Regardless, the presence of errors in the inference is unlikely to produce false positive PofO associations, since inference errors are expected to be drawn independently from the phenotypes. Instead, errors are expected to lead to false negatives as PofO signals get diluted onto the two parental origins and thus decrease association power. In this work, we controlled for this by focussing exclusively on high-confidence PofO calls, which corresponds to a call for 74.5% of heterozygous genotypes with an estimated error rate below 1%. The overall high accuracy in our estimates could be achieved thanks to recent progress in the statistical estimation of haplotypes for very large sample sizes^15,45^ so that the PofO status inferred within IBD tracks could be confidently propagated to entire chromosomes. Of note, we think that further improvements in phasing algorithms can be made by explicitly modelling IBD sharing between close relatives, eventually through inter- and intra-chromosomal scaffolding as we proposed in this work.

Our ability to infer PofO depends on the availability of close relatives. Surprisingly, even when only a single third degree relative is available for IBD mapping, we achieve a high call rate and a low error rate. We believe this could be further improved by using more distant relatives, even though they will not contribute as much as second- and third-degree relatives to the inference. In addition, our PofO inference depends on our ability to assign parental status to relatives based on IBD sharing on chromosome X, which comes with some flaws. First, our current inference is only possible for males as it leverages chromosome X haploidy, which means that only male specific PofO effects can be investigated. Potential improvements should come with whole genome sequencing (WGS) data: parental status assignment based on rare variant matching on chromosome Y and eventually mitochondrial DNA would likely become possible. In the UK Biobank, this has the potential to substantially increase the sample size above the ~26,000 samples we have so far to a theoretical upper bound of 105,826 samples, which corresponds to the number of samples for which we found groups of close relatives in the dataset. This could further boost the discovery power of downstream PofO association scans. Second, this approach can be confounded by high levels of inbreeding which could lead a sample to share portions of the chromosome X IBD with close relatives on both sides of the family, therefore greatly complexifying sex assignment. However, we consider this issue to be almost negligible in this study as the UK biobank mostly comprises outbred individuals.

Overall, this study is a valuable resource to further characterize PofO effects and catalogue imprinting genes in the human genome. We expect our approach to be applicable to other biobanks, such as those collected by the FinnGen research project (https://finnqen.qitbook.io/documentation/) or the Million Veteran Program^46^. Collective efforts would allow the detection of PofO effects with an unprecedented sample size by meta-analysing PofO effects across multiple biobanks and therefore greatly help future research on the molecular mechanisms leading to PofO effects and their implication for human health.

## Supplementary Figures/Tables Legends

**Figure S1:**
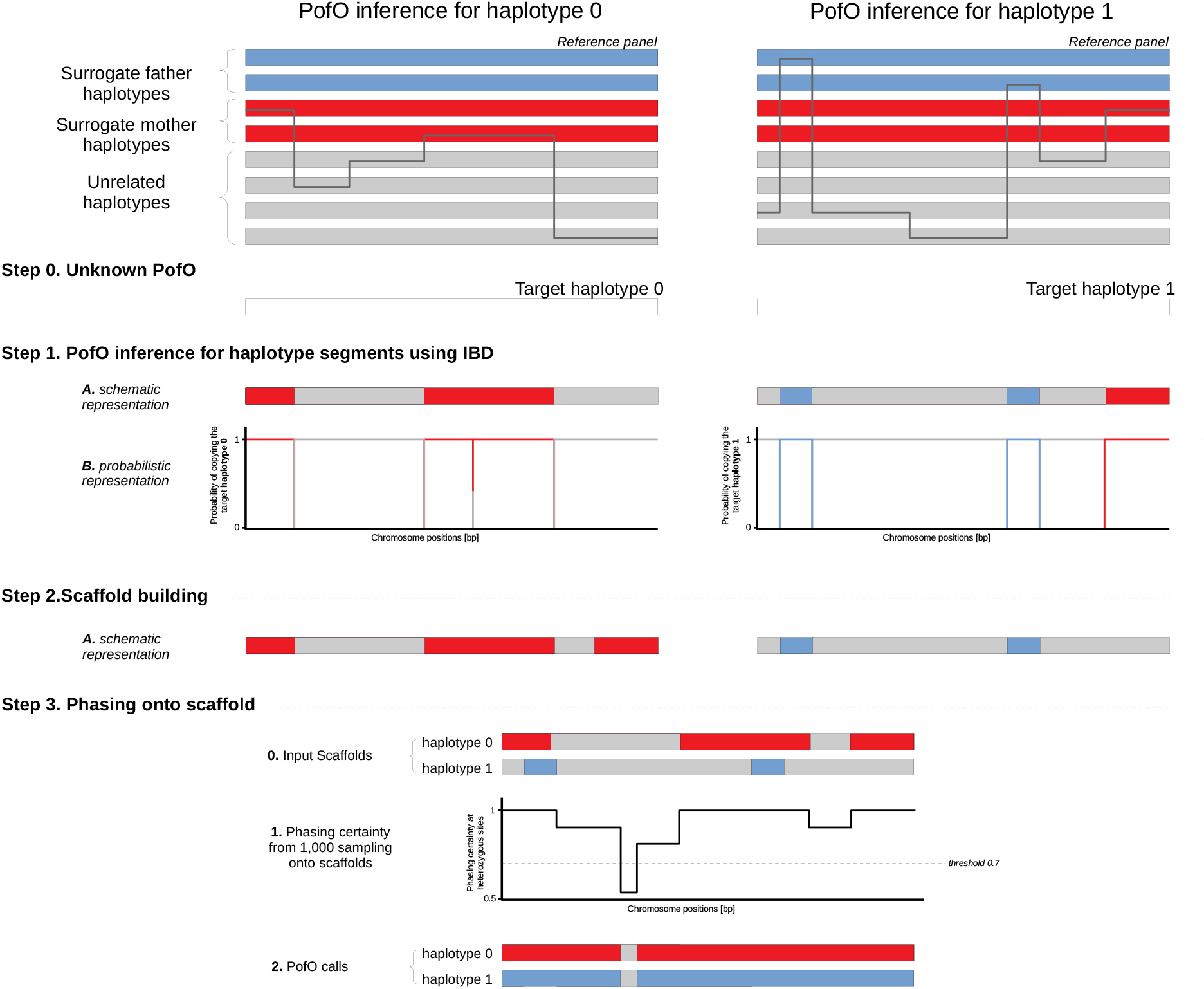
PofO inference from surrogate parents. Schematic representation of the PofO inference performed for each of the two haplotypes of a target sample. In step 1, the PofO of haplotype segments is inferred from the surrogate parents using a HMM that computes IBD sharing between the target haplotype and a reference panel comprising haplotypes from the surrogate parents and 100 unrelated haplotypes. In step 2, the resulting IBD segments are used to assemble a haplotype scaffold to inform another round of phasing. In step 3, we sampled a 1,000 haplotype pairs per target as part of the last phasing round in order to compute the frequency at which unassigned alleles co-localize onto the maternal/paternal scaffold, thereby deriving confidence scores for their PofO extrapolation.

**Figure S2:**
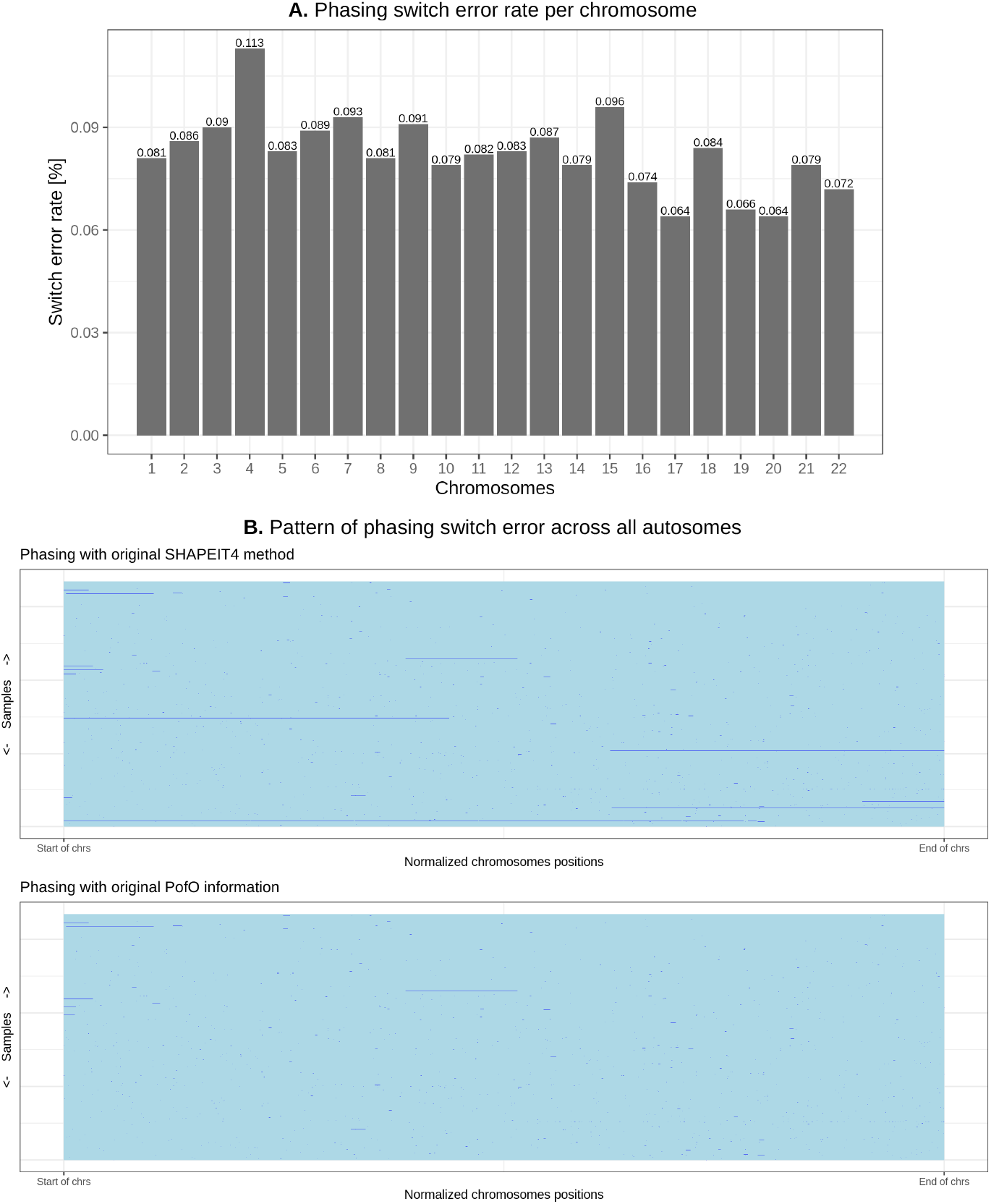
Phasing accuracy with known PofO information. **A.** Phasing switch error rates per chromosome. **B.** Genomic location of switch errors along the chromosomes for 308 samples with surrogate parents and genotyped parents (y-axis). Positions are normalized between 0 and 1 (x-axis). Continuous segments with the same colour represent segments correctly phased. Switches between two colours represent switch errors.

**Figure S3:**
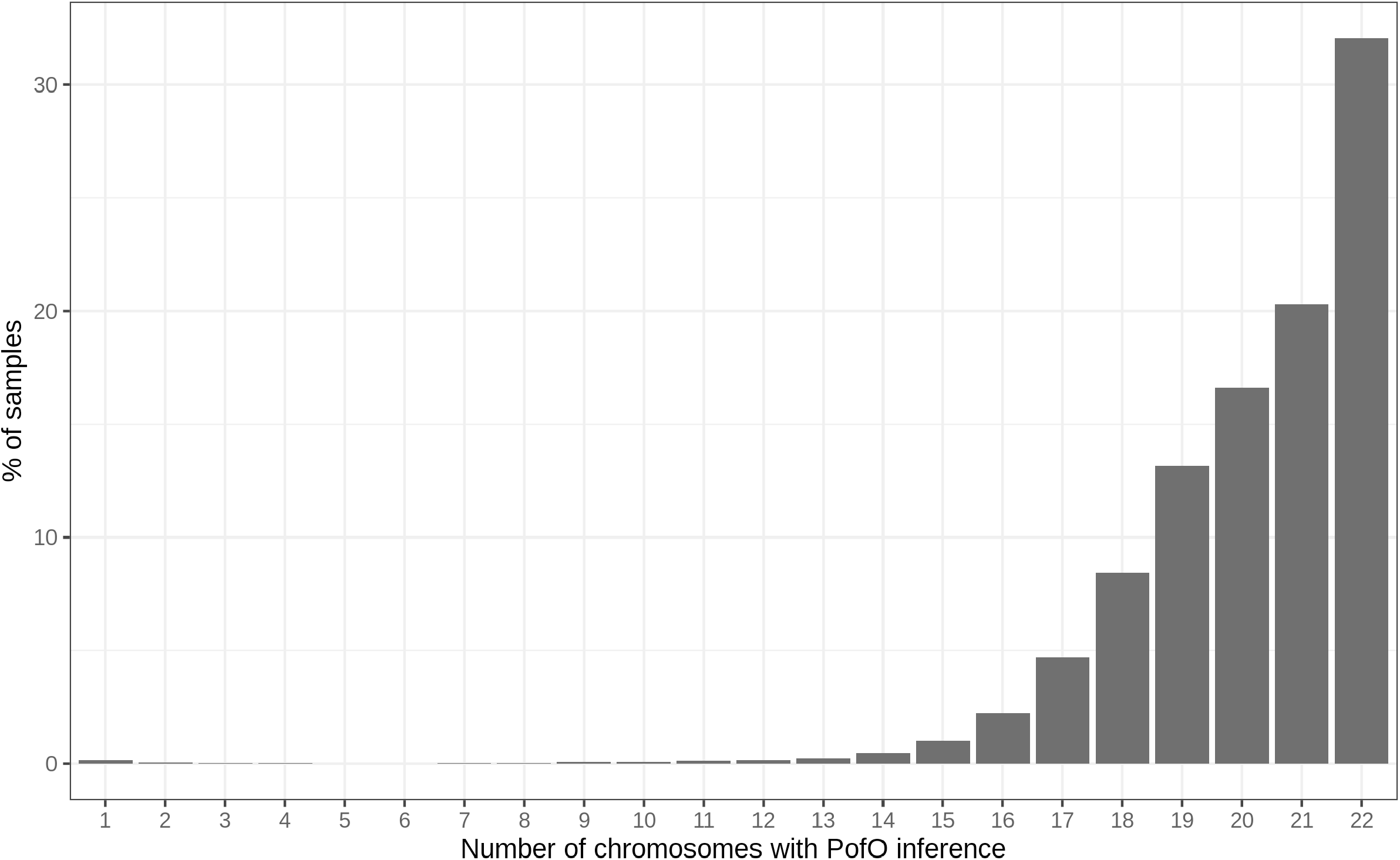
Coverage of PofO inference across samples. Proportion of samples (y-axis) having PofO inference across N chromosomes (x-axis).

**Figure S4:**
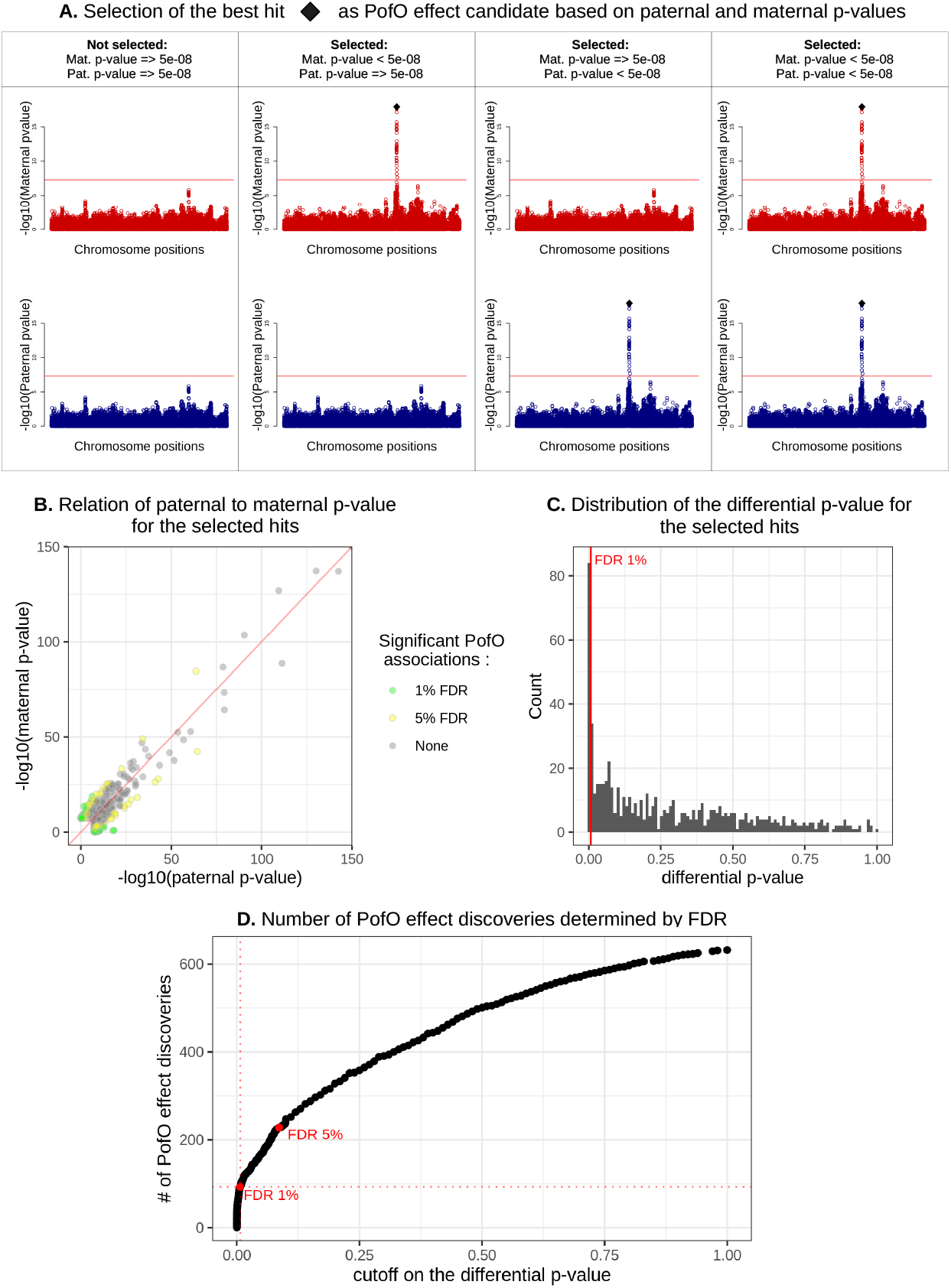
Selection of PofO associations. **A.** Schematic representation of the approach used to select candidate PofO effects. **B.** Scatter plot of the maternal and paternal p-values for the 631 candidate associations. Grey dots represent non-significant associations. Yellow and green dots represent associations being significant at 1% and 5% FDR, respectively. **C.** Distribution of the differential p-values across all 631 candidate PofO associations. **D.** Number of significant PofO associations (y-axis) as a function of the threshold used for differential p-values (x-axis). The red dots represent the 1% and 5% cut-offs on the differential p-value. A FDR of 1% has been used in this study, resulting in 92 significant associations.

**Figure S5:**
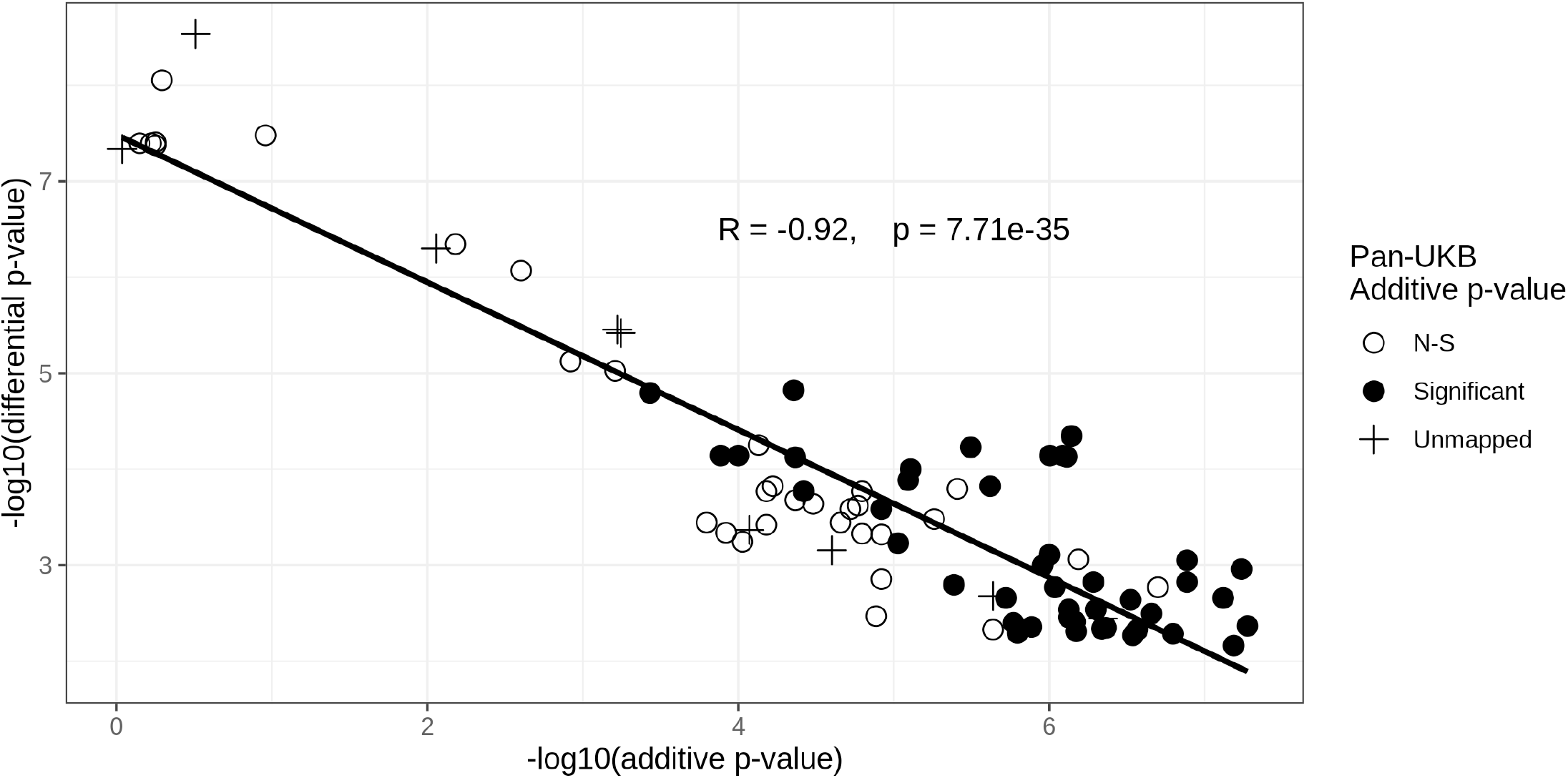
Relationship of non-additive PofO effect p-values. Scatter plot of the additive and differential p-values for all PofO associations non-significant in the additive scan. Dots are coloured in black or white depending if they are significant or not in the full UK Biobank, respectively (N=420,531 Europeans; Pan-UKB team; https://pan.ukbb.broadinstitute.org).

**Figure S6:**
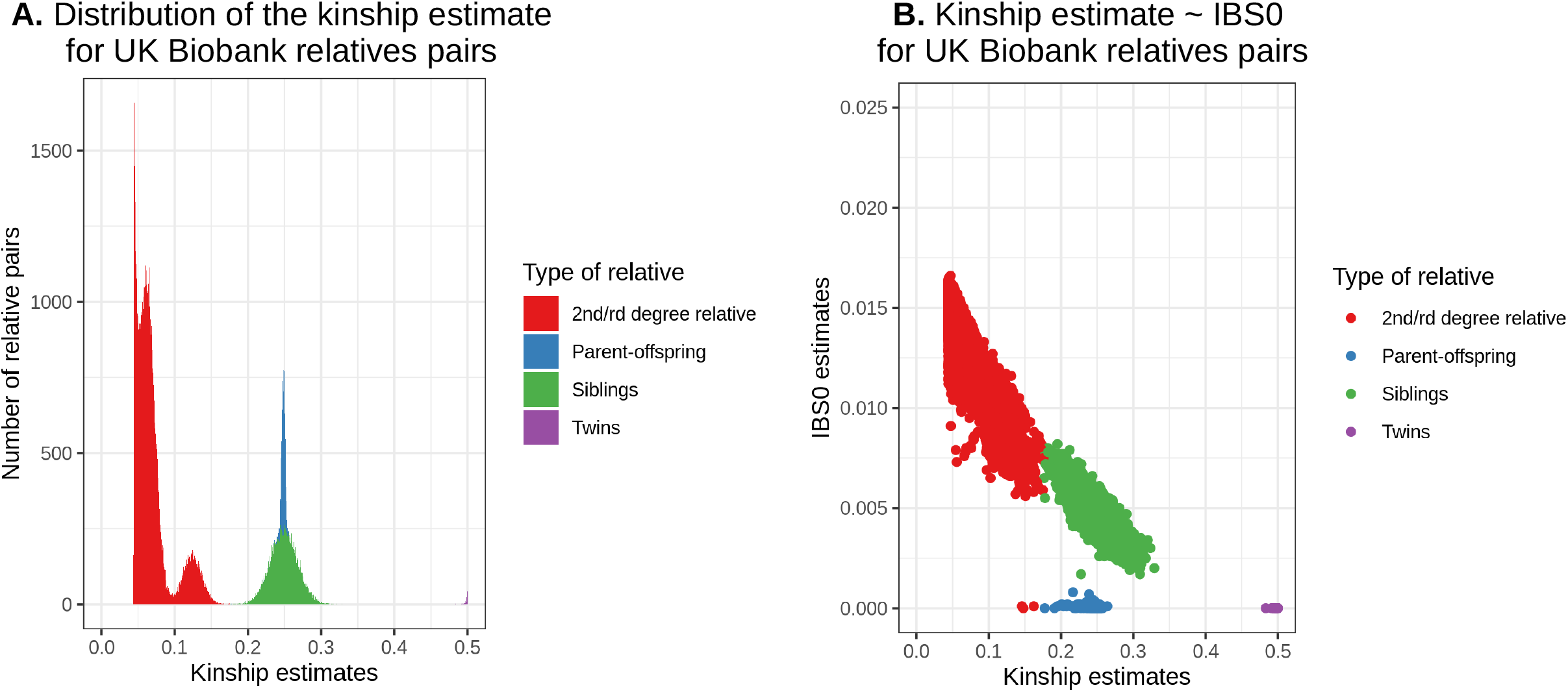
Relatedness in UK biobank. **A.** Distribution of the kinship estimate across all UK Biobank relative pairs up to the third degree. **B.** Kinship estimates in function of the IBS0 estimates in all UK biobank relative pairs up to the third degree. Colours indicate the classification used in our analysis.

**Figure S7:**
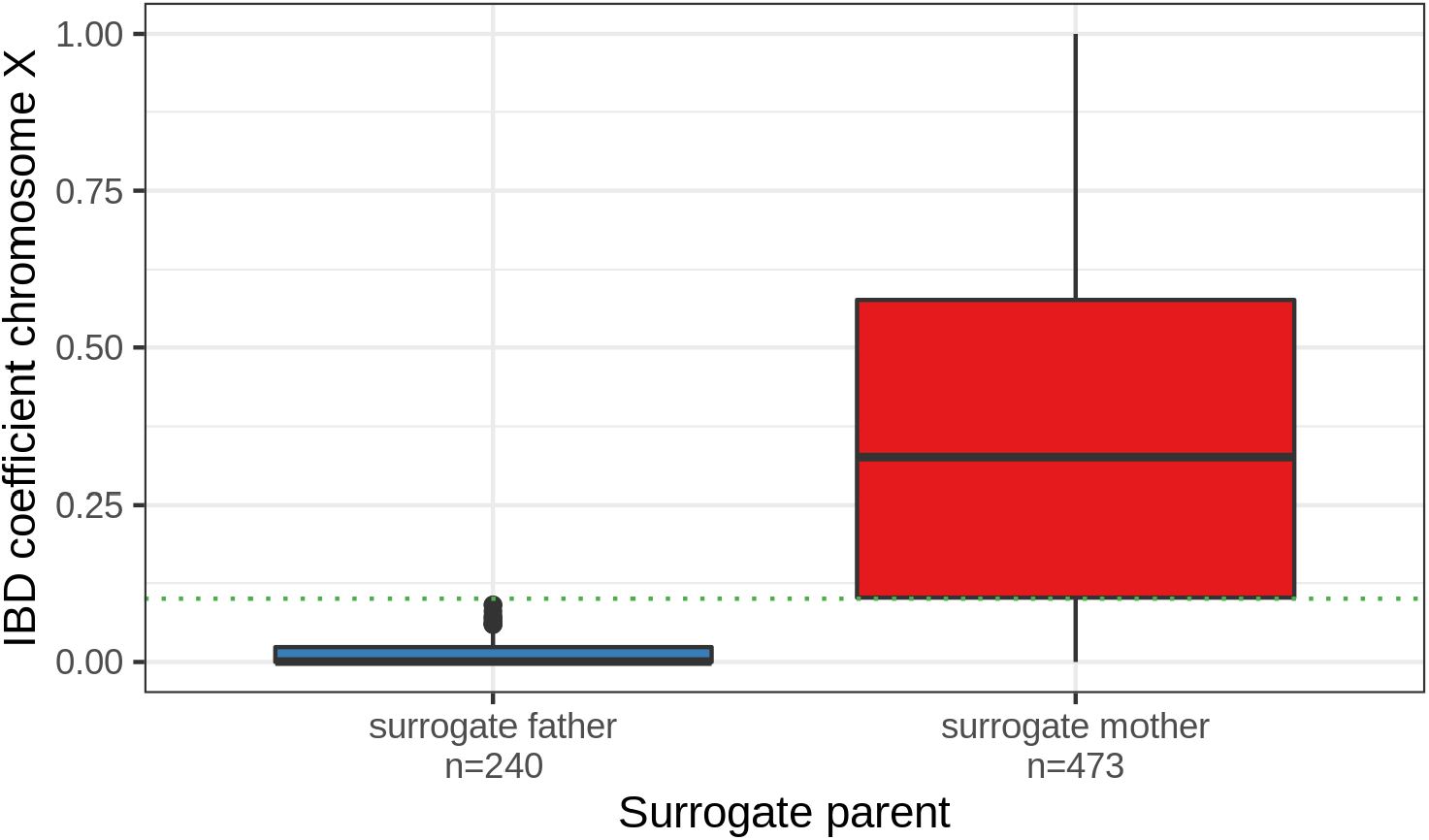
Validation of chromosome X IBD. IBD coefficient on chromosome X (y-axis) for targets with genotyped parents and surrogate parents (i.e. validation cohort). The IBD is estimated between these targets and their surrogate mother or surrogate father (x-axis). Dotted green line indicates the cut-off used in our analysis (=0.1).

**Figure S8.**
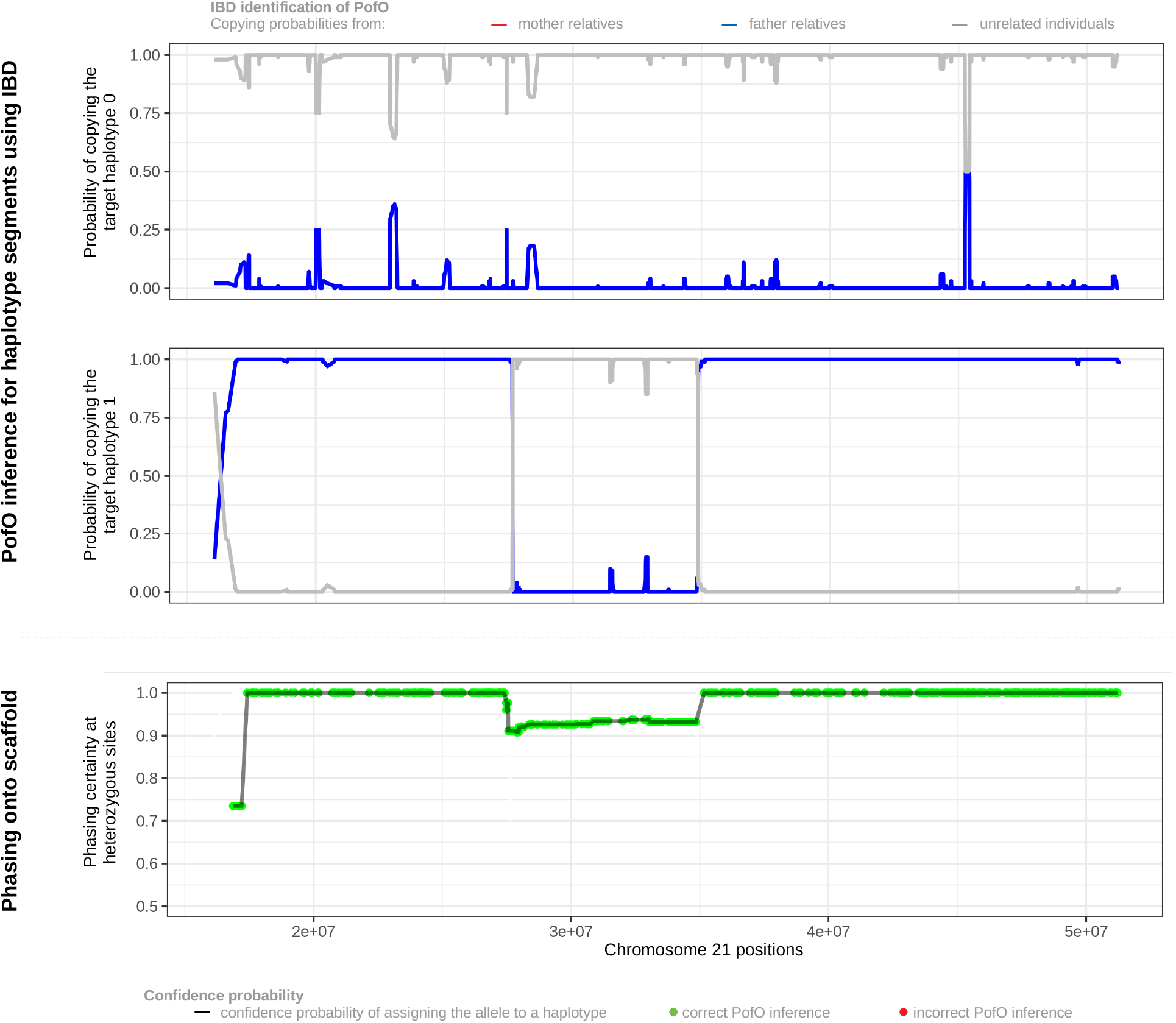
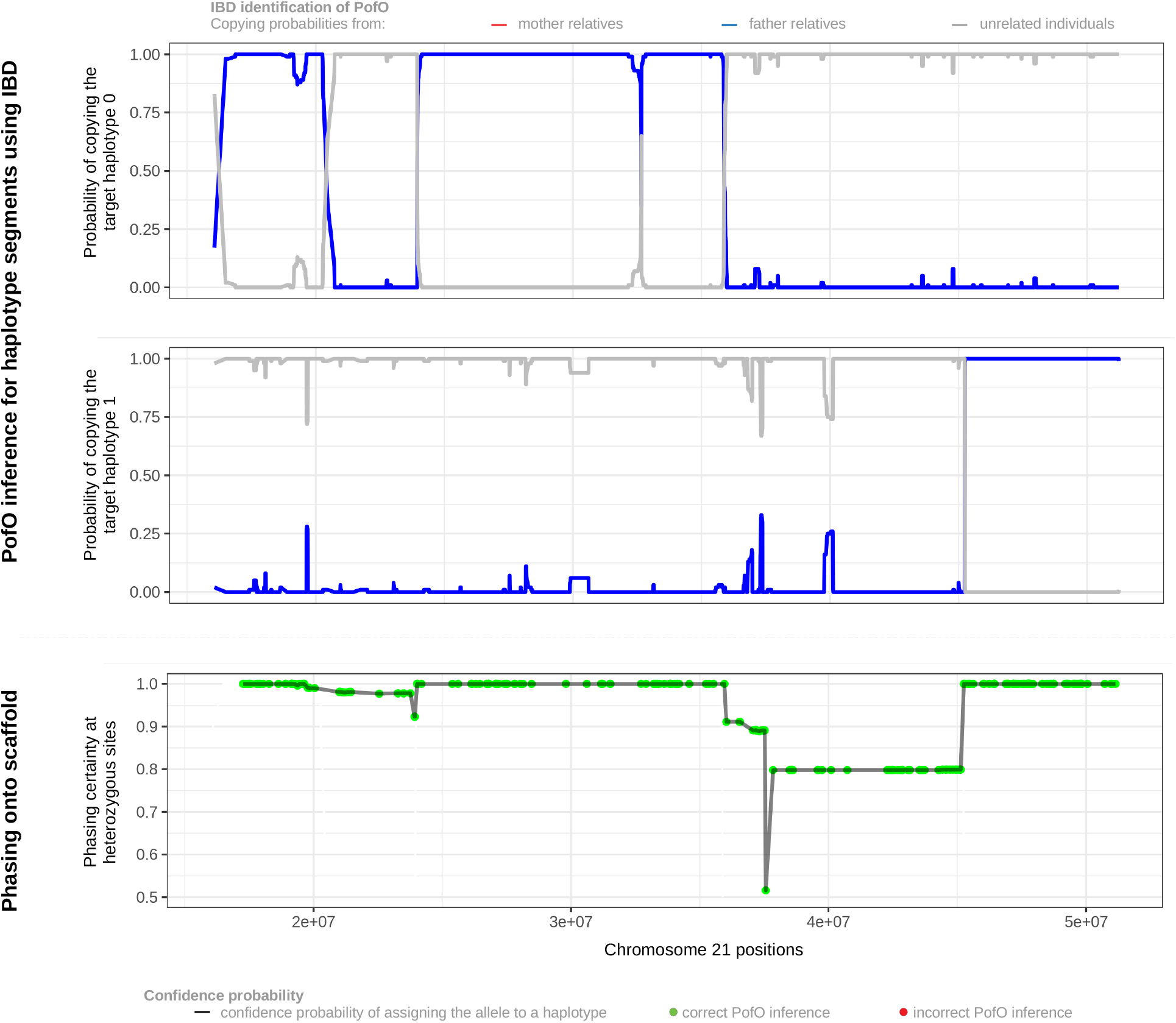
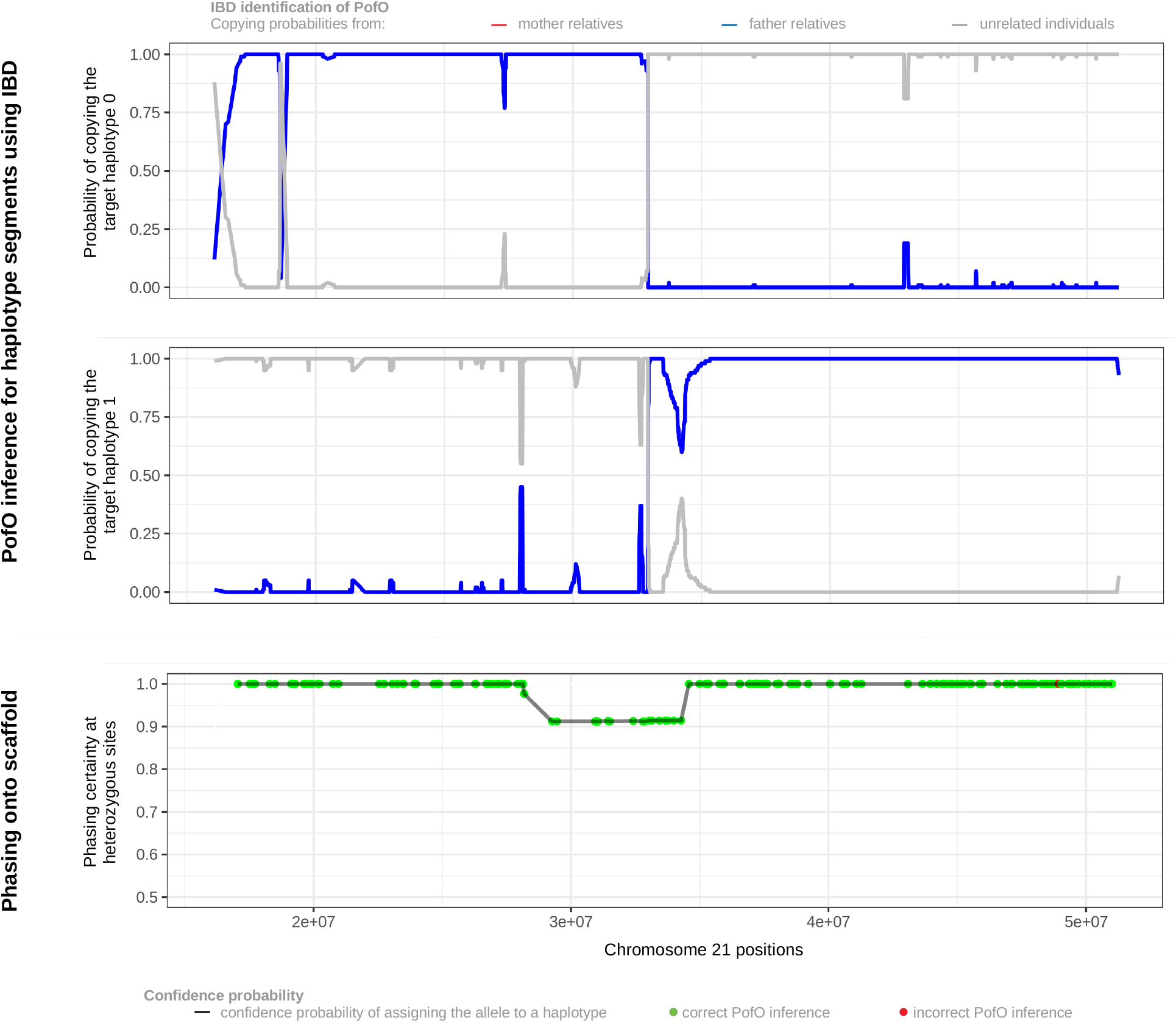
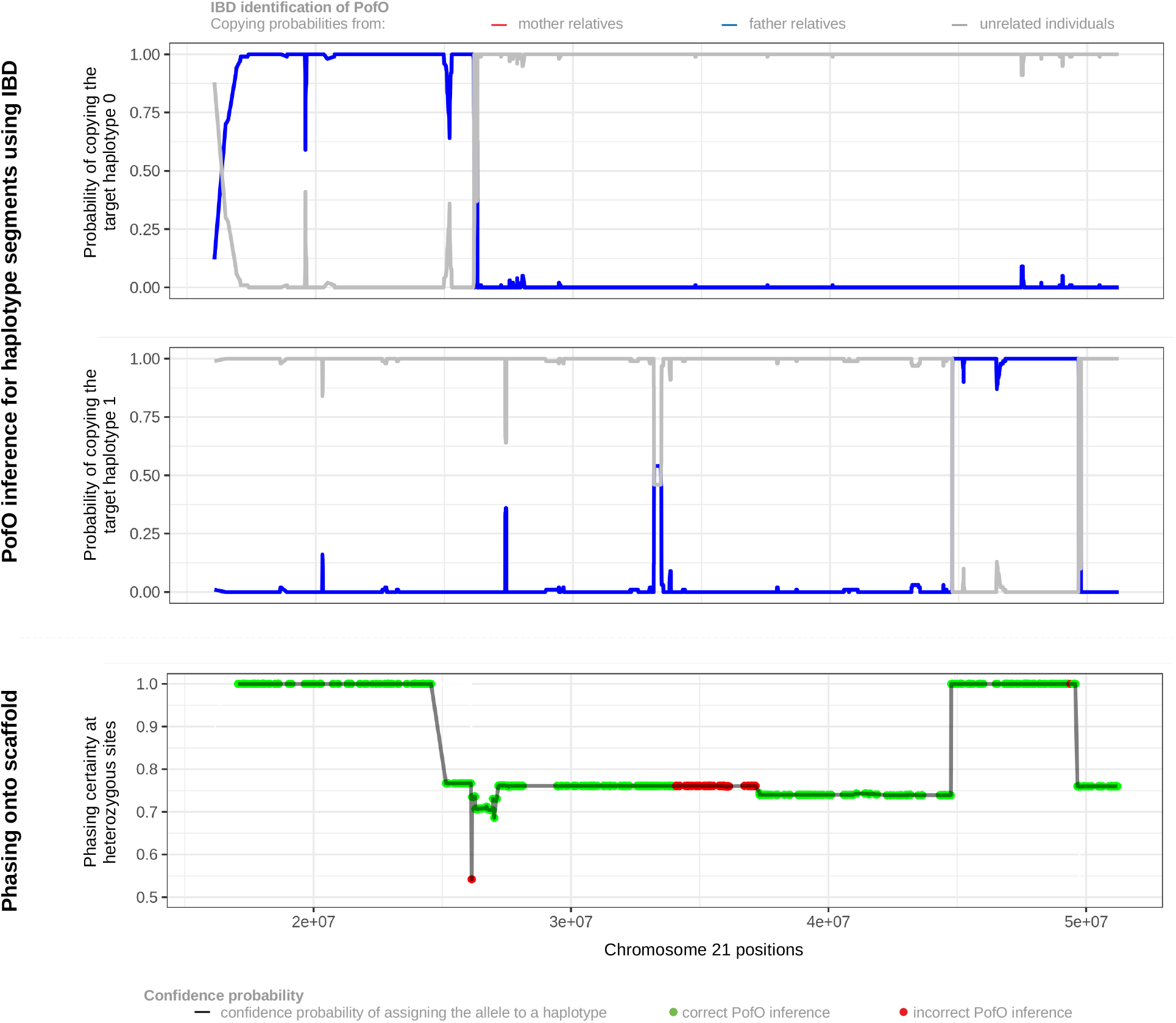
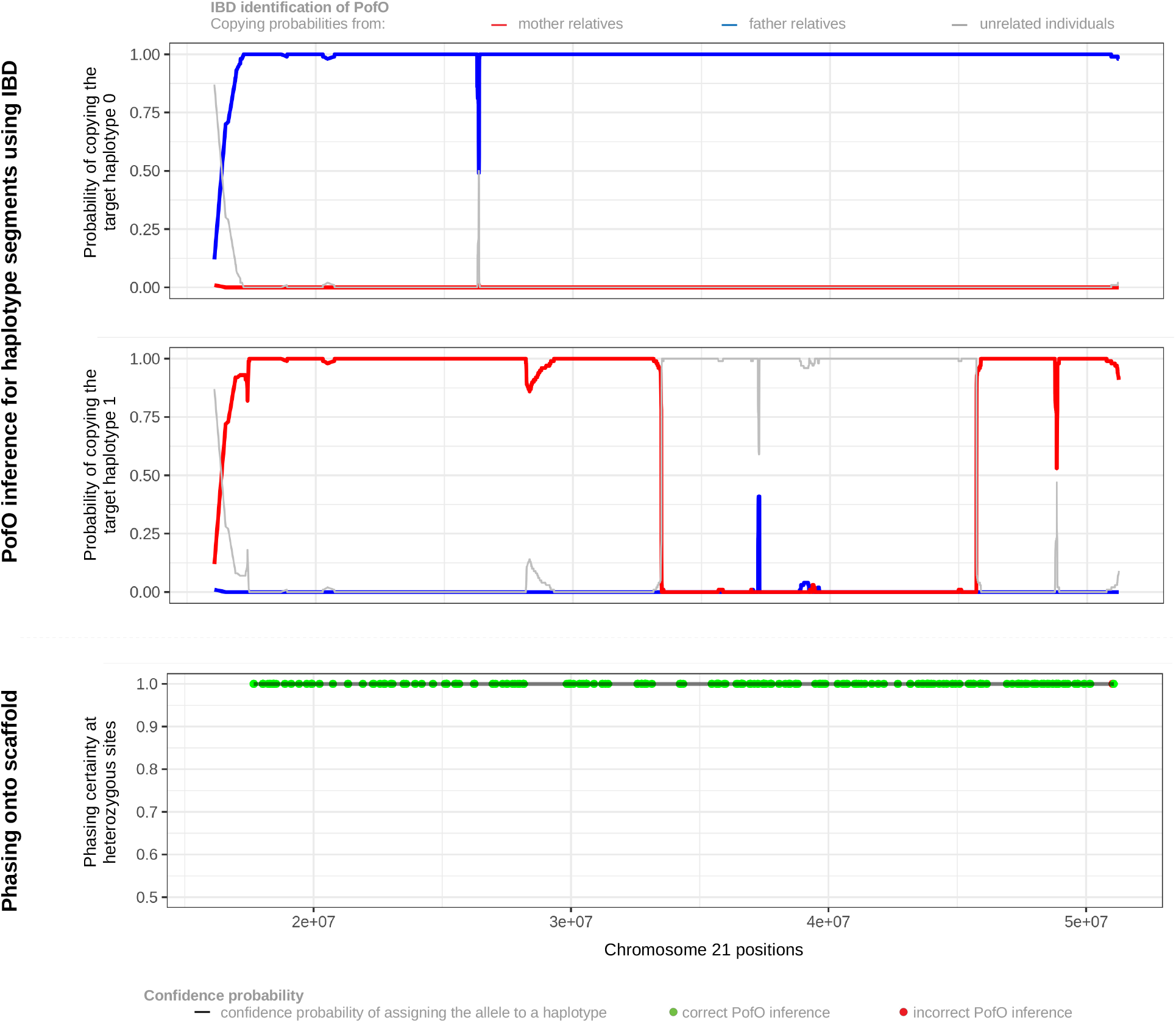
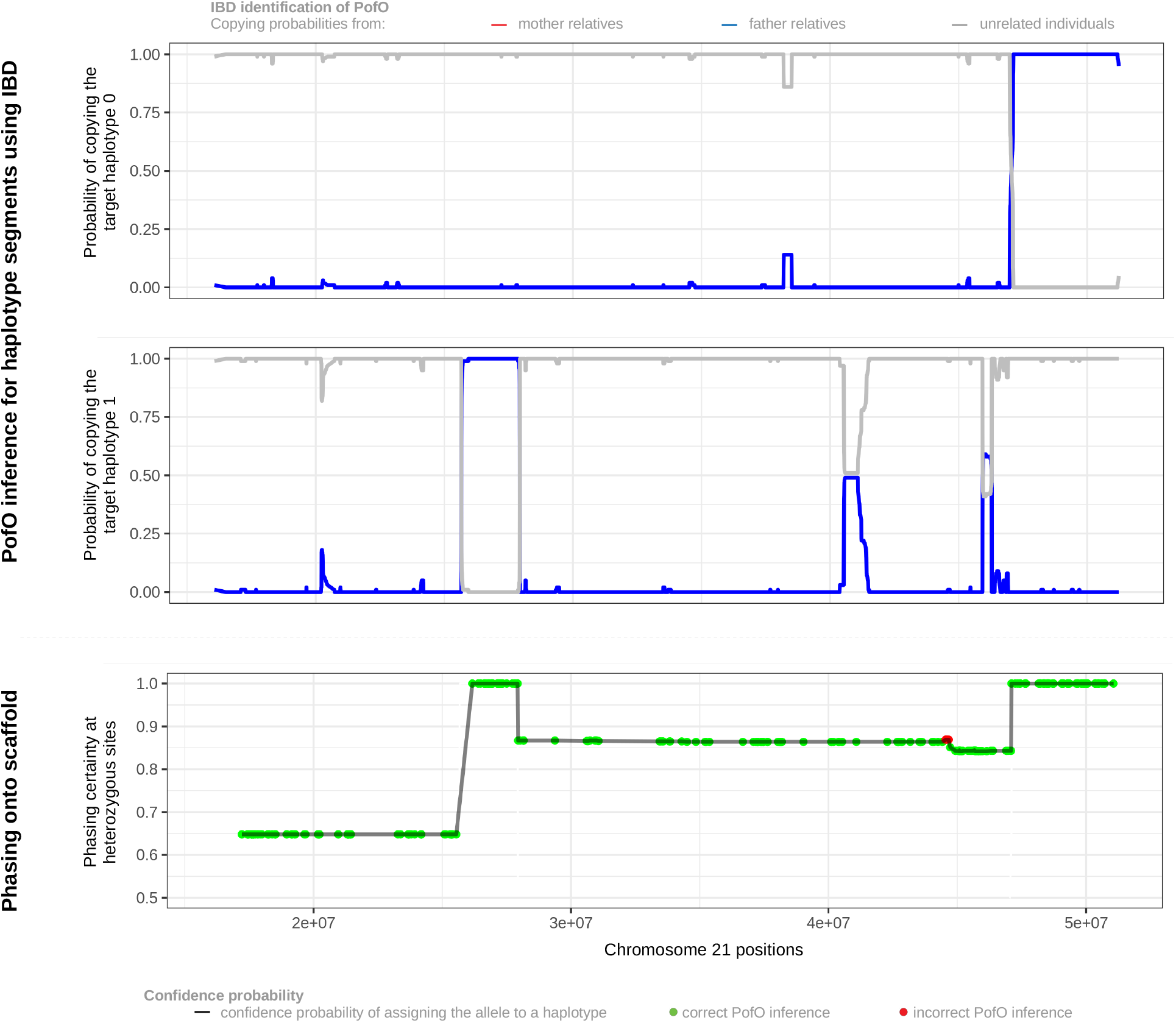
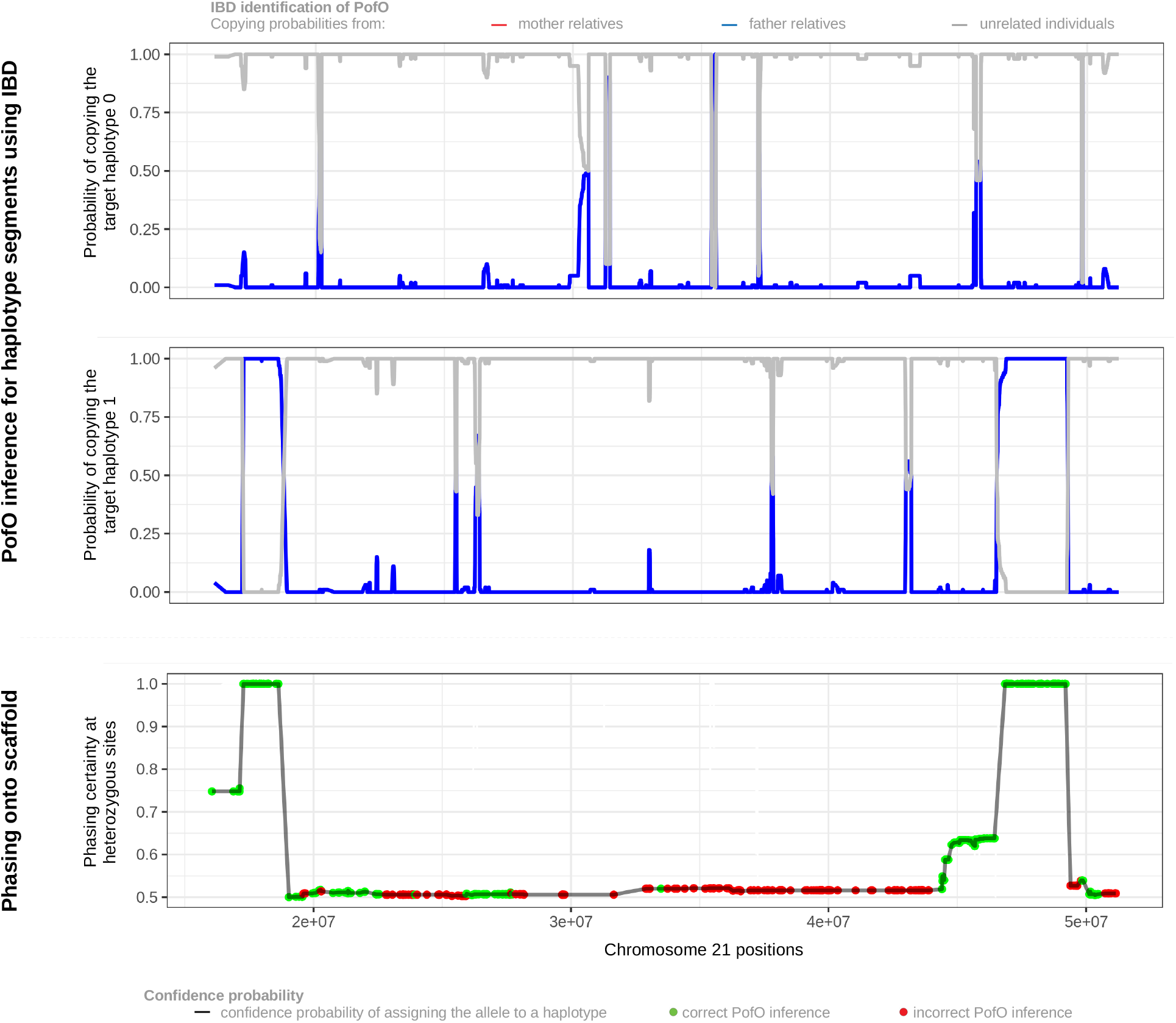
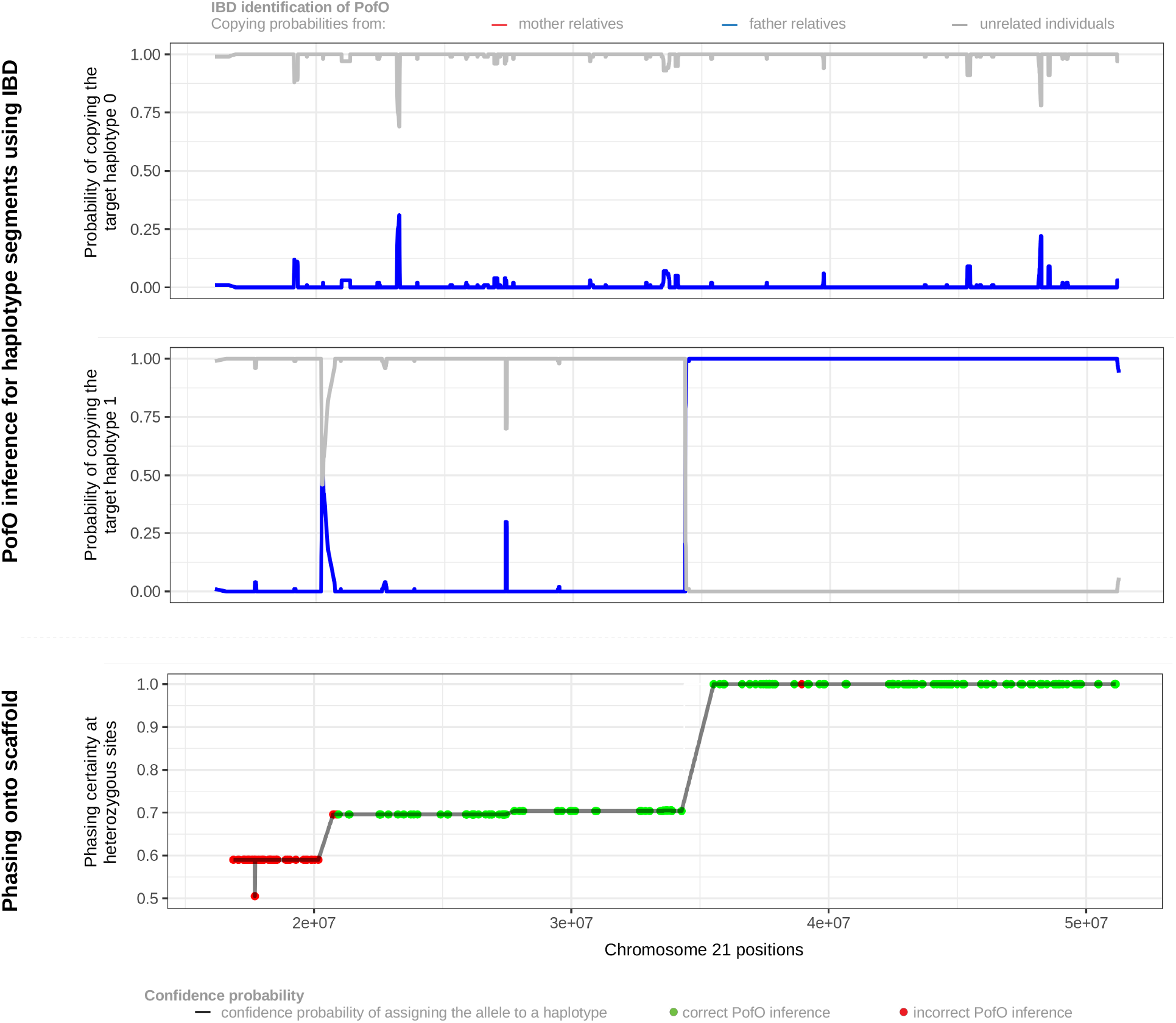
A-H. PofO inference from the surrogate parents for chromosome 21 across 8 validation individuals. Panels 1-2 show the probability given by our HMM (y-axis) that each target haplotype copies from the surrogate father(s) in blue, the surrogate mother(s) in red or unrelated individuals in grey as we move along chromosome 21. Panel 3 shows the phasing certainty resulting from 1,000 sampling onto scaffold. Red dots show errors in the PofO inference while green dots show correct PofO inference.

## Methods

### Duos/Trios identification

To identify trios and duos we used pairwise kinship and IBS0 estimates up to third degree relative computed using KING^13^ and provided as part of the UK biobank study. Following Manichaikul *et al.*^13^ (Table 1) and Bycroft *et al.*^12^ (Supplementary material), we defined offspring-parent pairs as having a kinship coefficient between 0.1767 and 0.3535 and an IBS0 below 0.0012 (**Figure S6**). We also added the condition of age difference greater than 15 years between parent-offspring pairs. We used the age and sex of the individuals to distinguish parents and offspring. For the trios, we also ensured that the two parents have different sex. Starting from 147,731 UKB individuals with at least one third degree relative, we found a total of (i) 1,064 samples with both mother and father (i.e. trios) and (ii) 4,123 samples with mother or father (i.e. duos). We used the reported ancestry of individuals to keep only genotyped individuals of British and Irish ancestry (N=443,993), which resulted in 1,037 trios and 3,872 duos.

### IBD based group inference

We used pairwise kinship and IBS0 estimates up to the third degree relative to identify sibling pairs (kinship between 0.1767 and 0.3535 and IBS0 above 0.0012), and second- and third-degree relatives’ pairs (kinship below 0.1767) for all genotyped individuals of British and Irish ancestry (N=443,993) (**Figure S6**). We found 106,414 individuals with at least one second or third degree relative and 21,255 sibling pairs. For individuals with two or more relatives, we separated those relatives into groups, representing the groups of relatives on each side of the family (i.e. mother-side relatives and father-side relatives). To do so, we used the relatedness in-between these relatives: those related to each other are expected to be on the same side of the family, while those unrelated to each other are expected to be on different sides of the family. We built for each individual a kinship symmetric matrix of size NxN, where N is the number of second-to-third relatives of the individual considered, filled with the kinship values in-between each relative. We then used the ‘igraph’ R package^47^ to cluster these relatives into groups based on their relatedness similarly to what has been done by Bycroft *et al.*^12^. As we wanted a maximum of two distinct groups (i.e. one paternal and one maternal), we excluded samples with more than two clusters of relatives from the analysis. We identified a total of 105,826 individuals with groups of relatives, ranging from one group of one relative to two groups of more than two relatives. This includes 309 individuals having also both parents genotyped (i.e. trios) and 1,090 having a single parent genotyped in the data (i.e. duos). These 1,399 individuals with at least one genotyped parent and groups of close relatives constitute our validation data set on which we applied our PofO inference method using the close relatives as surrogate parents, ignoring the parental genomes. We then used parental genomes to compute the accuracy of our inference.

### Groups assignments

We assigned parental status (i.e. mother or father) to groups of close relatives by examining shared IBD segments on chromosome X using XIBD^48^, a software specifically designed to map IBD on chromosome X (**Figure 1.C**). This assignment was only possible for males as they inherit their only chromosome X copy from their mother: a close relative sharing IBD on chromosome X with the target is expected to be from the maternal side of the family. To empirically determine the IBD threshold above which only mother-side relatives are found, we used the 1,399 samples of our validations set (i.e. with close relatives’ groups and genotyped parents). We computed the IBD sharing on chromosome X for each target-relative pair, knowing the correct parental side of the relatives from the kinship in between the relatives and the available parents. We found that only mother-side relatives share more than 0.1 of IBD1 on chromosome X (**Figure S7**), a value that we used as a threshold to assign maternal status. Across the 107,038 individuals having groups of close relatives, 48,814 individuals are males, and we assigned the group of close relatives to the maternal side of the family for 20,620 of them. By extension, we propagated the maternal status to the relatives from the same parental group, and we labelled as paternal the relatives from the other group. We then used the underlying idea that siblings share the same set of cousins, uncle and aunt to enrich our set of samples. We searched for siblings of these 20,620 individuals having the exact same close relatives’ groups. We found 864 such siblings, resulting in a total of 21,484 individuals with close relatives’ groups assigned to parental status (i.e. surrogate parents).

### Genotype processing

We used the UK biobank SNP array data in PLINK format. We converted the UK biobank plink files into VCF format using PLINK v1.90b 5^49^, which resulted in 784,256 variant sites across the autosomes for 488,377 individuals. We then used the UK biobank SNPs QC file (UK biobank resource 1955) to keep only variants used for the phasing of the original UK biobank release, resulting in 670,741 variant sites.

### Validation and production datasets

We assembled two distinct datasets comprising different collections of samples of British and Irish ancestries. The first one includes all UK Biobank samples excluding the parental genomes for the N=1,399 validation samples for which we have both parental genomes and surrogate parents. We ran our inference on N=1,399 validation samples and we assessed its performance by comparing our estimates to the truth given by parental genomes. It is important to note that parental genomes have been used only at the validation stage and not during any phasing runs nor PofO inference. The second dataset includes this time all available UK Biobank samples and has been used to produce the final set of individuals with PofO inference that has been used for association testing. This includes N=21,484 samples for which PofO could be inferred from surrogate parents and N=4,909 samples for which PofO could be directly inferred from the trios/duos.

### PofO inference step1: IBD mapping

In this first stage, we inferred PofO for alleles shared IBD with surrogate parents. To do so, we started by an initial phasing run of the data using SHAPEIT v4.2.1^15^ with default parameters so that all data consists of haplotypes. Then, we designed a Hidden Markov Model (HMM)^50^ to identify IBD sharing between the target haplotypes and a reference panel mixing haplotypes from 2 different sources: from the surrogate parents of the target (labelled as mother or father) and from unrelated samples. We aimed for such a probabilistic model for its robustness to phasing and genotyping errors compared to approaches based on exact matching such as the positional Burrows–Wheeler transform (PBWT). The model then uses a forward-backward procedure to compute, for each allele of a target haplotype, the probability of copying the allele from (i) the surrogate mother haplotypes, (ii) the surrogate father haplotypes or (iii) unrelated haplotypes. Here, we used 100 unrelated haplotypes as decoys so that the model is not forced to systematically copy from surrogate parents. When the model copies the target haplotype from a specific surrogate parent at a given locus with high probability, we can therefore infer the PofO at this locus from the parental group the surrogate parent belongs to. When the model copies from unrelated haplotypes, no inference can be made at the locus (**Figure S1, Figure S8 panel 1,2**). We implemented this approach in an open-source tool available on GitHub (https://github.com/RJHFMSTR/PofO_inference). As a result of this procedure, we obtained PofO calls within haplotypes segments shared IBD with surrogate parents.

### PofO inference step2: extrapolation by phasing

In this second stage, we inferred PofO for all remaining genotyped alleles. First, we built a haplotype scaffold comprising all alleles for which we know PofO from IBD sharing with surrogate parents^14^. In other words, we forced all alleles that we knew to be co-inherited from the same ancestor to locate on the same homologous chromosome (**Figure S1, Figure S2B**). In the scaffolds, we only included IBD tracks longer than 3cM. We empirically determined this length on the validation set of samples by maximizing and minimizing the call rate and the error rate, respectively (see method section *Accuracy and parameters optimization).* In addition, we considered in the haplotype scaffold only alleles having a PofO probability greater or equal to 95%. As a result of this, we could build paternal and maternal haplotype scaffolds that we used in a second step to rephase the entire dataset using SHAPEIT4 v4.2.1^15^. The goal of this second round of phasing is three-folds: (i) to ensure that the pool of alleles coming from the same parent land onto the same haplotype, (ii) to propagate the PofO assignment from IBD tracks to all alleles along the chromosomes and (ii) to correct long range switch errors. Point (ii) is made possible as all alleles with PofO unknown (i.e. not in IBD tracks) are phased relatively to the haplotype scaffold so that we can extrapolate their PofO from the scaffold they co-localize with (paternal/maternal). In practice, we ran SHAPEIT4 with two main options: *--scaffold* to specify the scaffolds of haplotypes to be used in the estimation and *--bingraph* to output the haplotype reconstructions together with phasing uncertainties. The latter provides the haplotype reconstructions as parsimonious graphs encapsulating phasing uncertainty so that likely haplotype pairs can be rapidly sampled without being forced to rerun the complete phasing run. As a consequence, we sampled for each target sample a 1,000 haplotype pairs using different seeds and computed the probability for a given allele to be paternal or maternal from its frequency of co-localization across the 1,000 pairs onto the paternal and maternal haplotype scaffolds, respectively **(Figure S1, Figure S8A-H panels 3)**. This frequency indicates the certainty we have in phasing and therefore is a probabilistic measurement of the confidence in the PofO assignment. For instance, a specific allele being phased with a certainty of 0.8 onto the paternal haplotype scaffold has an 80% chance to be of paternal origin. In all downstream analysis, we considered only heterozygous genotypes with a phasing probability above 0.7; a threshold that we empirically determined from the validation set of samples by maximizing and minimizing the call rate and the error rate (see method section on *Accuracy and parameters optimization*).

### PofO inference step3: extrapolation by imputation

In this third stage, we inferred PofO for untyped alleles, i.e. not included on the SNP array. To do so, we imputed the data using IMPUTE5 v1.1.4^16^ with the Haplotype Reference Consortium^17^ as a reference panel. As our data is phased with each haplotype assigned to a specific parent, we used the parameter *--out-ap-field* to run a haploid imputation of the data and separately imputed the paternal haplotype and the maternal haplotype. Of note, we filtered out all heterozygous genotypes with a phasing certainty below 0.7 prior to imputation (see previous section). As a result of haploid imputation, the PofO of imputed alleles can be probabilistically deduced from the imputation dosages: an allele imputed with a dosage of 0.85 on the paternal haplotype has 85% probability of being inherited from the father (i.e. PofO probability = 85%). Finally, we filtered out variants with an INFO score below 0.8 and obtained a dataset comprising 22,156,064 variants.

### Accuracy and parameters optimization

We used samples with both genotyped parents and groups of surrogate parents (i.e. validation set of samples N=1,399) to compute the errors in the PofO inference and to optimize the parameters of our inference method. For the trios (N=309) and the duos (N=1,090), we determined the correct parental origin of offspring heterozygous genotypes at sites where a parent is homozygous, excluding sites with Mendel inconsistencies. We assessed the impact of two parameters on the call rate (percentage of heterozygous genotypes with PofO assignment) and the error rate (percentage of heterozygous genotypes with incorrect PofO assignment) of the PofO inference: (i) the length in centimorgan (cM) of the haplotype segments that we included in the scaffold for the second phasing run and (ii) the phasing certainty threshold we used to assume PofO to be known at heterozygous genotypes. To do so, we compute the call rate and the error rate for all combinations of the following parameters (**Figure 2A**): haplotype segments of 2cM, 3cM, 5cM, 8cM and 10cM and threshold on the phasing certainty between 0.5 and 1.0 by steps of 0.05. Overall, we found that a phasing certainty above 0.7 and haplotype segments above 3cM to be a good trade-off between call rate and error rate, and used these values in all downstream analyses.

### Association testing for PofO

We tested 97 quantitative phenotypes of the UK biobank data set **(Table S1**) from 4 phenotypic categories: body size measurements, body composition by impedance, blood biochemistry and blood count. From these categories, we considered only those with at least 50% of the UK Biobank samples being phenotyped. We rank-transformed each phenotype using the ‘rntransform’ function from the GenABEL v1.8-0 R package^51^. We used the sex, age and the method used to infer the PofO of alleles as covariates (i.e. surrogate parents or direct parents). We used BOLT-LMM v2.3.4^18^ to run all association tests. As recommended by the authors, we performed the model fitting only on the genotyped variants. For the additive GWAS scans, we used the *--dosageFile* parameter to test imputed alleles dosages, as recommended in the documentation. For the PofO GWAS scans (i.e. maternal scan and paternal scan), we used the *--dosageFile* parameter to test the PofO dosages of alleles. In practice, we only used imputed allele dosages (i) of the paternal haplotype for the paternal-specific GWAS and (ii) of the maternal haplotype for the maternal-specific GWAS, so that PofO assignment uncertainty is propagated to association testing. We conducted a third PofO GWAS scan that compares the effect of maternally and paternally inherited minor alleles at heterozygous genotypes (i.e. differential scan). For this, we used only heterozygous genotypes with imputed minor allele dosages greater or equal to 0.95 to keep only genotypes with high confidence in the PofO. We encoded such alleles as 0 when inherited from the father and 1 when inherited from the mother. We again used the *--dosageFile* parameter to test whether the paternal and maternal alleles have differential effect at heterozygous sites. All homozygous genotypes have been set to missing. Prior to running association testing, we re-ordered all alleles so that we systematically tested the effects of minor alleles. We filtered out all variants with a minor allele frequency (MAF) below 5% which resulted in 5,426,121 variants for association testing.

### GWAS hits identification

We used the Bonferroni whole genome significance to identify significant independent associations (p<5e^-8^) for each phenotype. We identify independent hits as having Linkage Disequilibrium (LD) < 0.05 and being located at least 500kb apart. If two hits are not independent, we select the one with the lowest p-value. LD is computed with PLINK v1.90b5^49^. We identified additive hits as having additive p-value < 5e^-8^. To identify PofO significant associations, we used a two-step approach. In the first step, we selected all associations genome-wide significant (Bonferroni; p-value < 5×10^-8^) in the differential association scan, resulting in 9 associations with strong differences in parental allele effect sizes. In the second step, we defined a set of putative PofO associations by selecting all those being genome-wide significant in the maternal scan, in the paternal scan or in both scans. From this set, we distinguished PofO effects from additive effects by keeping only associations significant at 1% FDR in the differential scan, resulting in 92 associations with strict paternal or maternal effects. FDR was computed using the R/qvalue package.

### URLs

GeneImprint, http://www.geneimprint.com/; Catalog of Imprinted Genes, http://www.otago.ac.nz/IGC.

## Acknowledgements

This work was conducted under UK Biobank project 66995 and funded by the Swiss National Science Foundation (SNSF) project grant 373 (PP00P3_176977).

## Author contributions

R.J.H. and O.D. designed the study and wrote the paper. R.J.H. performed experiments. R.J.H. and O.D. developed the IBD mapping algorithm. S.R. helped with the imputation. D.M.R helped with biological interpretations. Z.K. helped with the design of the GWAS models. This study was initiated after discussions between A.B. and O.D. The project has been supervised by O.D. All authors reviewed the final manuscript.

## Data availability

The summary statistics for the four GWAS models across the 97 phenotypes are available here for download: http://poedb.dcsr.unil.ch/

## Code availability

Repository https://github.com/RJHFMSTR/PofO_inference hosts the source code of the IBD mapper used as part of this study.

## Supplementary tables

https://docs.google.com/spreadsheets/d/1bof9TFsqSCGXSuZH_irPeuYgioEPztyrvcvfCnTCkwo/edit?usp=sharing

